# Characterizing the molecular regulation of inhibitory immune checkpoints with multi-modal single-cell screens

**DOI:** 10.1101/2020.06.28.175596

**Authors:** Efthymia Papalexi, Eleni Mimitou, Andrew W. Butler, Samantha Foster, Bernadette Bracken, William M. Mauck, Hans-Hermann Wessels, Bertrand Z. Yeung, Peter Smibert, Rahul Satija

## Abstract

The expression of inhibitory immune checkpoint molecules such as *PD-L1* is frequently observed in human cancers and can lead to the suppression of T cell-mediated immune responses. Here we apply ECCITE-seq, a technology which combines pooled CRISPR screens with single-cell mRNA and surface protein measurements, to explore the molecular networks that regulate *PD-L1* expression. We also develop a computational framework, *mixscape*, that substantially improves the signal-to-noise ratio in single-cell perturbation screens by identifying and removing confounding sources of variation. Applying these tools, we identify and validate regulators of *PD-L1*, and leverage our multi-modal data to identify both transcriptional and post-transcriptional modes of regulation. In particular, we discover that the kelch-like protein *KEAP1* and the transcriptional activator *NRF2*, mediate levels of *PD-L1* upregulation after IFNγ stimulation. Our results identify a novel mechanism for the regulation of immune checkpoints and present a powerful analytical framework for the analysis of multi-modal single-cell perturbation screens.

## INTRODUCTION

Immune checkpoint (IC) molecules regulate the critical balance between activation and inhibition during immune responses. Under normal physiological conditions, inhibitory IC molecules are essential to maintain self-tolerance and prevent autoimmunity [1,2], but their expression is often mis-regulated in human cancers to escape immune surveillance [3,4]. For example, the inhibitory IC *PD-L1*, which interacts with the *PD-1* receptor on T cells to inhibit T-cell activation [5], is overexpressed in many cancers and serves as a prognostic factor for patient survival and response to immunotherapy [6]. There is therefore substantial interest not only in identifying therapeutic avenues to block these interactions, but also in understanding the molecular networks utilized by cancer cells to up-regulate ICs like *PD-L1*.

Previous efforts have established an initial set of molecular regulators that influence both mRNA and surface protein levels for *PD-L1*. Numerous studies have observed that exposure to interferon gamma (IFNγ) rapidly induces *PD-L1* expression both in cancer cell lines *in vitro*, as well as in the tumor microenvironment [7–10]. Core components of the IFNγ response therefore represent upstream regulators of *PD-L1* expression, including the transcription factor IRF1 (which binds directly to the *PD-L1* promoter [11]), the *JAK-STAT* signal transduction pathway, and the IFNγ receptors themselves. Additional modulators of IFNγ signaling [12], *PD-L1* promoter chromatin state [13], or response to UV-mediated stress [14] have also been identified. In addition, there has been particular recent interest in the characterization of putative post-transcriptional regulators of *PD-L1* stability and degradation. For example, the Cullin 3-*SPOP* E3-ligase complex can directly ubiquitinate PD-L1 in a cell-cycle dependent manner, leading to its degradation [15]. In addition, a genome-wide CRISPR screen identified two previously uncharacterized regulators, CMTM6 and CMTM4, which stabilize PD-L1 surface expression by preventing lysosome-mediated degradation [16,17]. In each of these cases, perturbation of *PD-L1* regulators was shown to modulate the activity of anti-tumor T cells, highlighting the therapeutic interest in understanding the regulation of inhibitory IC molecules.

We recently introduced expanded CRISPR-compatible CITE-seq (ECCITE-seq), which simultaneously measures transcriptomes, surface protein levels, and perturbations at single-cell resolution, as a new approach to identify and characterize molecular regulators [18]. ECCITE-seq builds upon the experimental design of pooled CRISPR screens, where multiple perturbations are multiplexed together in a single experiment, but offers distinct advantages. First, the single-cell sequencing readout (i.e. Perturb-seq, CROP-seq, CRISP-seq) [19–21], enables the measurement of detailed molecular phenotypes, instead of one phenotype (expression of a single protein or cell viability). Second, by simultaneously coupling measurements of mRNA, surface protein, and direct detection of guides within the same cell [22], ECCITE-seq allows for multimodal characterization of each perturbation. We therefore reasoned that ECCITE-seq would enable us to simultaneously test and identify new regulators of IC molecules, and in particular, to distinguish between transcriptional and post-transcriptional modes. Moreover, the rich and high-dimensional readouts readily facilitate network and pathway-based analyses, which could go beyond the identification of individual genes and yield insights into their regulatory mechanism.

Here, we apply ECCITE-seq to simultaneously perturb and characterize putative regulators of inhibitory IC molecules in response to IFNγ stimulation. When analyzing our single-cell data, we identified confounding sources of heterogeneity, including the presence of cells that received a targeting guide RNA but exhibited no perturbation effects, introducing substantial noise into downstream analyses. We developed and validated computational methods to control for these factors, and substantially increased our statistical power to characterize multi-modal perturbations.

Leveraging these tools, we identify a set of genes whose perturbation affects *PD-L1* transcript levels, surface protein levels, or both, and characterize the underlying molecular pathways utilized by each regulator. In particular, we find that the kelch-like protein *KEAP1* and the transcriptional activator *NRF2*, both of which are frequently mutated in human cancers [23], can modify *PD-L1* levels. We link these findings to a novel regulatory mechanism for *CUL*3, and show that this gene acts as an indirect transcriptional activator of *PD-L1* mRNA via stabilization of the *NRF2* pathway. Taken together, our findings identify an important pathway for immune checkpoint regulation, and present a powerful and broadly applicable analytical framework for analyzing ECCITE-seq data.

## RESULTS

Human cancer cells routinely up-regulate IC molecules, such as *PD-L1*, to escape immune surveillance. The blockade of these checkpoints can significantly enhance the efficacy of the anti-tumor immune response, particularly during immunotherapy [24]. We were therefore motivated to gain deeper understanding of the molecular pathways and regulators that affect inhibitory IC expression, with a particular focus on *PD-L1*. Aiming to develop an experimental system to study multiple ICs simultaneously, we screened four cancer cell lines (THP-1, K562, KG-1 and U937) and tested their ability to up-regulate IC molecules in response to cytokines by flow cytometry (Supplementary Methods). We found that stimulating THP-1 cells with a combination of IFNγ, Decitabine (DAC), and transforming growth-factor beta 1 (TGFβ1) resulted in robust induction of three ICs: PD-L1, PD-L2, and CD86 (Supplementary Figure 1A). We also created a modified THP-1 cell line to inducibly express Cas9 under doxycycline treatment, representing an in-vitro model system amenable to environmental and genomic perturbations (Supplementary Methods).

**Figure 1.**
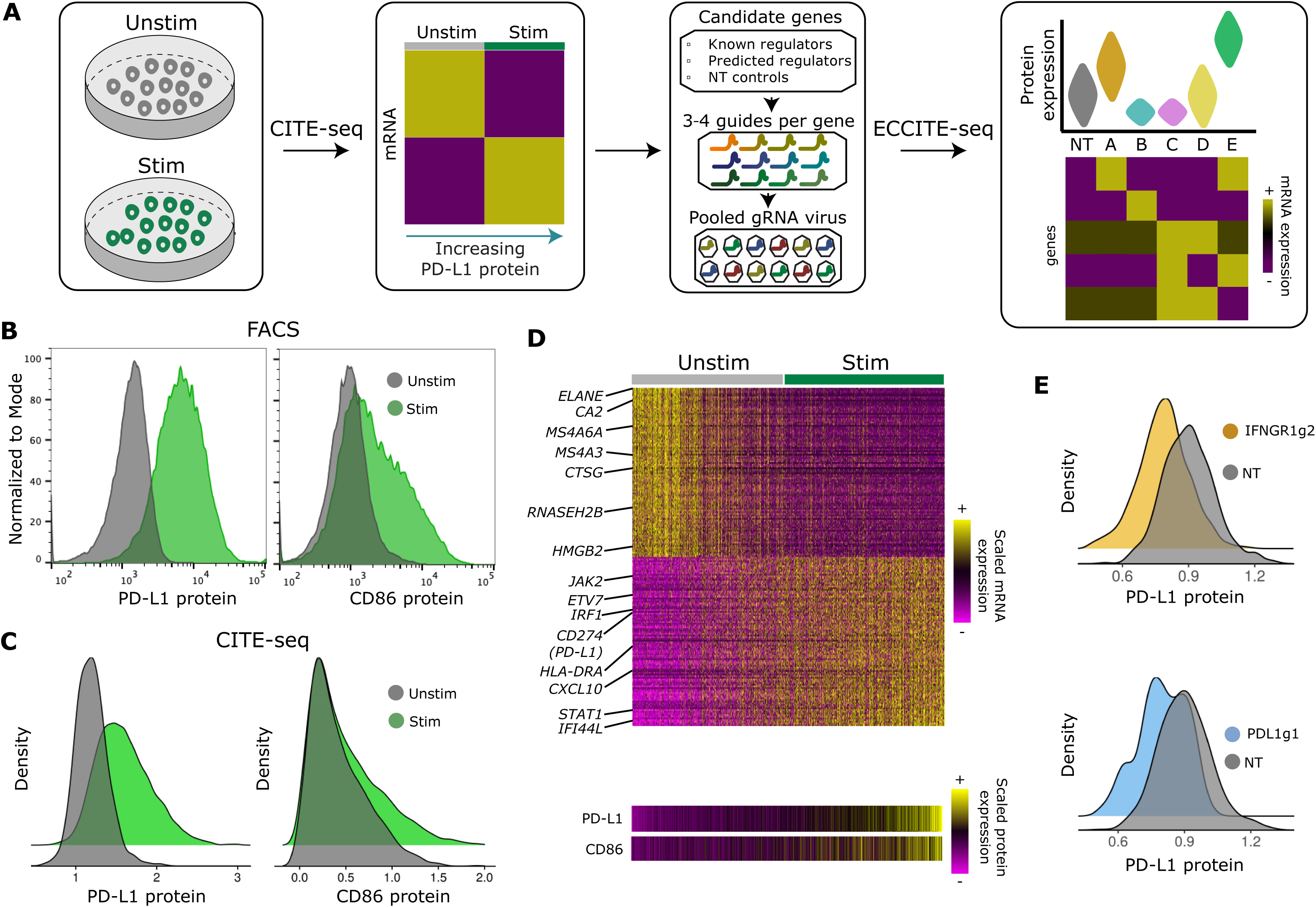
CITE-seq and ECCITE-seq identify regulators of PD-L1 protein expression. **(A)** Experimental design schematic. **(B)** Expression of PD-L1 (left) and CD86 (right) protein in stimulated (green) and control (grey) THP-1 cells, as measured by flow cytometry and **(C)** CITE-seq. **(D)** Single-cell heatmap showing the z-scored expression of 200 genes whose expression correlates with CD86 and PD-L1 protein expression (Supplementary Methods). **(E)** ECCITE-seq measurements of PD-L1 protein expression in cells that received gRNA targeting *PD-L1* and *IFNGR1*, and non-targeting controls.

In order to identify and characterize new regulators, we pursued a two-step experimental strategy, where each step leveraged multi-modal single-cell sequencing technologies (Figure 1A). First, we performed CITE-seq [22] on both unstimulated and stimulated THP-1 cells. Through the use of DNA-barcoded antibodies, CITE-seq enables the simultaneous measurement of cellular transcriptomes alongside surface protein levels of PD-L1, PD-L2, and CD86. We reasoned that these data would enable us to identify gene modules whose transcriptional levels mirrored the surface expression of each IC. Within these modules, we could identify a ‘target set’ of putative regulators representing genes known to affect transcription, chromatin, signaling, or protein stability. In a second step, we performed multiplexed perturbation and functional characterization of our target set. To accomplish this, we applied our recently developed ECCITE-seq technology, which extends CRISPR-compatibility to the CITE-seq protocol and enables simultaneous guide RNA capture. ECCITE-seq allowed us to multiplex >100 individual perturbations together, and to simultaneously test the effect of each in a single experiment. Moreover, the rich and multi-modal nature of these data allowed us to distinguish both transcriptional and post-transcriptional effects, and to explore mechanistic hypotheses for each gene.

### CITE-seq and ECCITE-seq enable identification and characterization of putative IC regulators

To identify putative IC regulators, we performed CITE-seq experiments on both stimulated and unstimulated THP-1 cells (Supplementary Methods). We recovered a total of 7,566 single-cell profiles, each representing coupled measurements of cellular transcriptomes and surface levels for three proteins: PD-L1, PD-L2 and CD86. For each surface protein, we compared the patterns of up-regulation upon stimulation observed by CITE-seq with those observed by flow cytometry, and found highly concordant results across technologies (Figure 1B, C; Supplementary Figure 1A, B). The multi-modal CITE-seq measurements allowed for the identification of genes whose expression is activated alongside IC surface protein induction (Supplementary Methods). Induced genes included well-characterized members of the IFNγ pathway (including the receptors *JAK2, STAT1*, and *IRF1*), while down-regulated genes (*ELANE, MS4A6A, CTSG*) were consistent with the monocyte progenitor identity of resting THP-1 cells.

Based on these results, we selected 26 genes for downstream characterization (Supplementary Table 1). Our panel included eight genes with well-characterized regulatory effects on *PD-L1*, and 18 genes representing transcription factors, chromatin regulators, signaling regulators, and modifiers of protein stability, that were mined from our CITE-seq data but where a clear link with IC regulation has not been firmly established. The first set represents positive controls for downstream analyses, while the second are putative new regulators. We designed a pooled single guide RNA (sgRNA) library consisting of three to four gRNAs per gene along with ten non-targeting (NT) guides, representing a total library of 111 gRNAs.

In order to functionally characterize our previously identified genes, we performed ECCITE-seq, a 5’ capture-based scRNA-seq method that is able to reverse transcribe sgRNA via the addition of a scaffold-specific primer, alongside cellular transcriptomes and ADTs. To guide our experimental design, we first performed a pilot experiment using gRNAs targeting *PD-L1* or *IFNGR1* as well as NT controls. In both cases, we observed a substantial reduction in PD-L1 expression, and perturbation of *IFNGR1* also ablated the IFNγ transcriptional response (Figure 1E). Clear effects were observed even after downsampling the dataset to 25 cells/gRNA (Supplementary Figure 1C).

We next performed an ECCITE-seq experiment utilizing our full library of 111 guides. Our total dataset represents three independent transductions (biological replicates) at low multiplicity of infection (MOI), aiming to maximize the proportion of cells infected with a single gRNA. After transduction, Cas9 expression was activated with doxycycline, and 90% of cells were stimulated to induce IC expression (the remainder were profiled without stimulation, Supplementary Figure 2A). Cells were then incubated with TotalSeq C antibodies (BioLegend), and processed on the 10x Genomics Single Cell 5’ assay. All samples were processed in parallel using our previously described multiplexing approach (‘cell hashing’; [25]), and sequenced on the Illumina NovaSeq platform (55,300 average mRNA reads/cell). Out of 30,328 cells, we found 22,606 cells where we could detect robust expression of at least one gRNA, including 22,573 where a cell could be specifically assigned to an individual perturbation (Supplementary Figures 2B-D), in line with the results of our pilot experiment.

**Figure 2.**
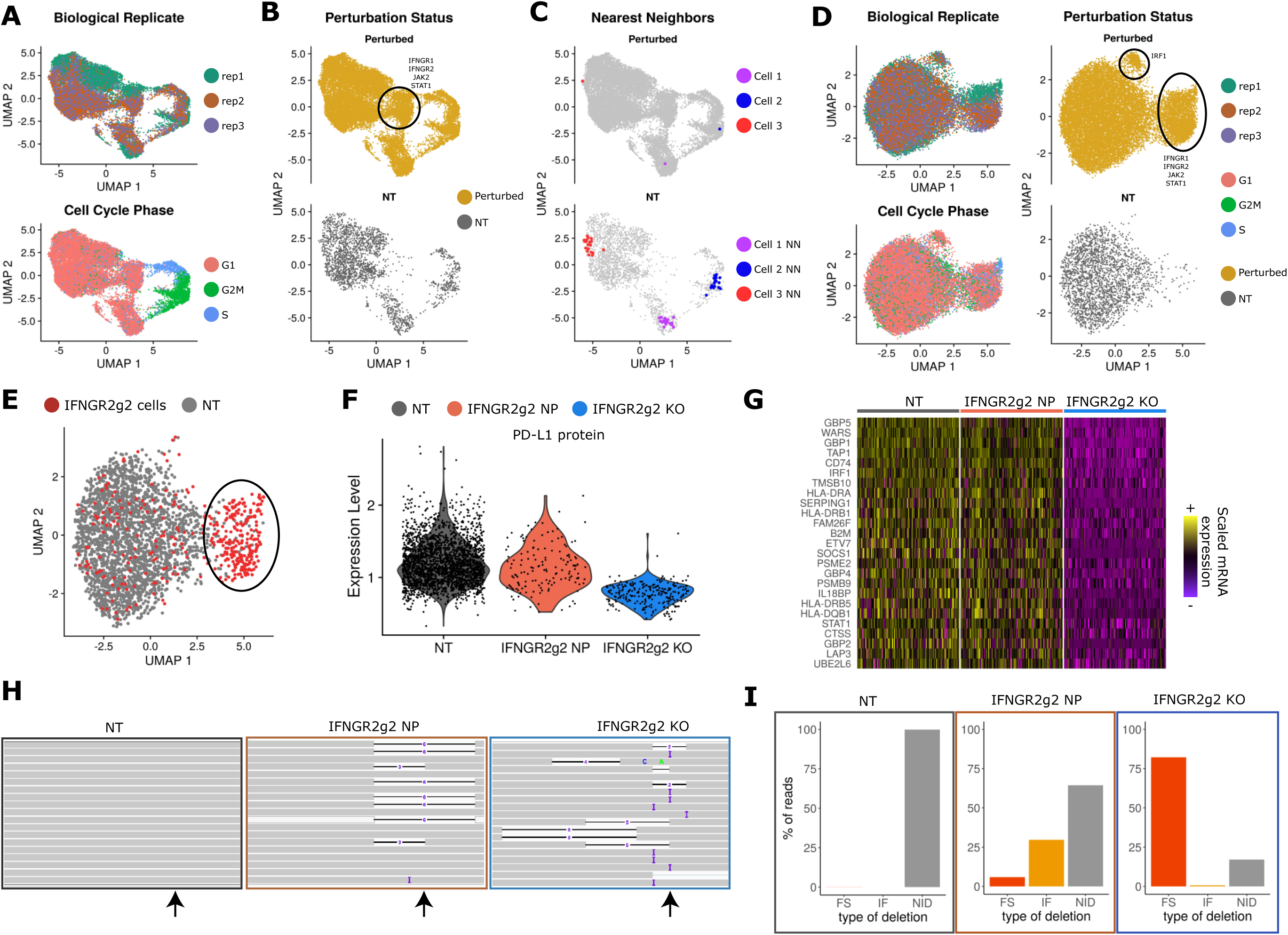
Calculating local perturbation scores removes unwanted sources of variation. **(A)** UMAP visualization of the ECCITE-seq dataset based on cellular transcriptomes. Cells are colored by biological replicate and cell cycle state. **(B)** Same as in A. Cells are split and colored by their perturbation status. Black circle denotes a perturbation-specific cluster. **(C)** Same as in B. Top panel: example of three distinct cells expressing an IRF1 gRNA (red, blue, purple). Bottom panel: their 20 nearest NT cell neighbors. Grey dots represent all remaining cells in the dataset. **(D)** UMAP visualization based on cellular perturbation scores. Black circles denote perturbation-specific clusters. **(E)** UMAP visualization showing all IFNGR2g2 and NT cells. Black oval denotes a group of putative IFNGR2g2 knockout (KO) cells that cluster separately, but a subset of targeted cells (outside the oval) appear to be non-perturbed (NP). **(F)** Violin plot showing PD-L1 protein expression in NT, NP, and KO cells. IFNGR2g2 KO cells exhibit low PD-L1 protein levels while IFNGR2g2 NP and NT cells express PD-L1 at identical levels. **(G)** Single-cell heatmap showing the mRNA expression of IFNγ pathway related genes in NT, NP, and KO cells. Gene expression is scaled (z-scored) across all single cells. For visualization purposes we downsampled our dataset to include 150 cells from each class shown in the heatmap. **(H)** Interactive Genome Viewer (IGV) screenshot of a representative sample of reads mapping at the *IFNGR2* gene locus (chr21: 34787276-34787299) targeted by IFNGR2g2 gRNA. CRISPR-induced insertions and deletions (INDELs) are seen in reads as black lines (I = insertion). gRNA cut site is denoted with a black arrow. **(I)** Barplot showing the % of reads with no INDELs (NID), inframe (IF) and frameshift (FS) mutations across NT, NP and KO cells. Only reads that overlapped the predicted cut site of IFNGR2g2 gRNA were used.

### Calculating local perturbation signatures removes confounding sources of variation

We next performed unsupervised dimensionality reduction (PCA) and visualization (UMAP) of the ECCITE-seq data based on their RNA profiles (Figure 2A, B; Supplementary Methods). While we had expected that cells would form groupings that were consistent with their underlying genetic perturbation, we initially observed that alternative sources of variation, including replicate identity, cell-cycle stage, and the activation of cellular stress responses (Supplementary Figure 3A, B), confounded our analysis. These sources of heterogeneity were also present in an independent analysis of NT control cells (those expressing non-targeting gRNAs, Supplementary Figure 3C), and we therefore designed a procedure to mitigate their effects.

**Figure 3.**
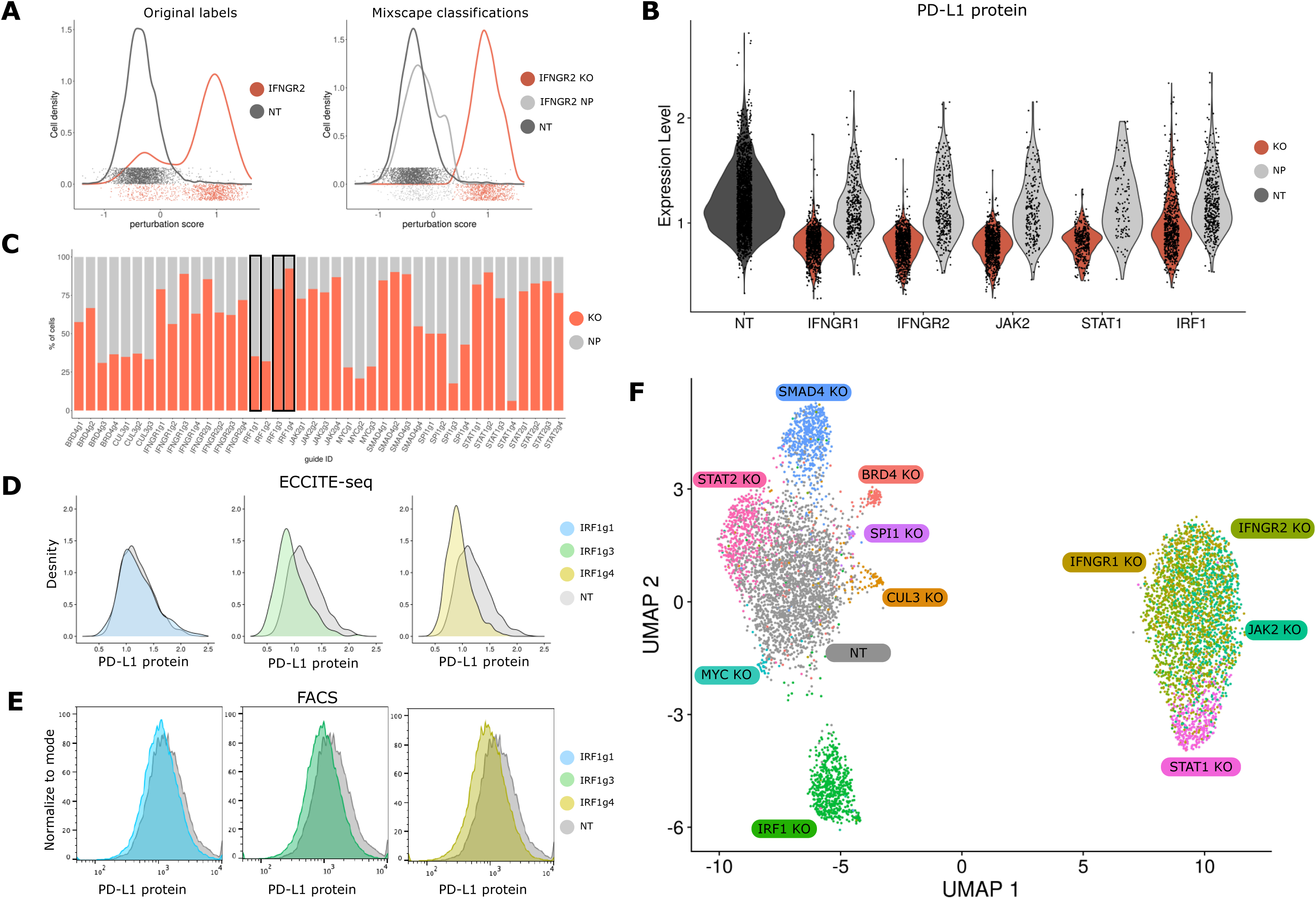
Mixscape removes cells that escape perturbation. **(A)** Distribution of perturbation scores (Supplementary Methods) for NT (grey) and IFNGR2 (red) cells. IFNGR2 cells are a mixture of two Gaussian distributions reflecting NP and KO cells. Classifying cells with mixscape resolves this heterogeneity. **(B)** Violin plot showing PD-L1 protein expression based on mixscape classification. Only KO cells show a reduction in PD-L1 protein levels when compared to NT control cells. **(C)** Barplot showing the percentage of targeted cells classified as KO by mixscape for each gRNA. Black box highlights three gRNAs targeting *IRF1* gene locus. **(D)** ECCITE-seq measurements of PD-L1 protein expression for cells expressing three distinct gRNA targeting *IRF1*, and NT controls. **(E)** Flow cytometry measurements of PD-L1 protein expression for the same populations as in (D). **(F)** UMAP visualization of all 7,421 NT and KO cells after running Linear Discriminant Analysis (LDA) (Supplementary Methods), revealing perturbation-specific clustering.

Briefly, for each target cell (expressing one target gRNA), we identified 20 cells from the control pool (NT cells) with the most similar mRNA expression profiles (Figure 2C; Supplementary Methods). These k=20 nearest neighbors should be in a matched biological state to the target cell, but did not receive a targeting gRNA. Therefore, subtracting their averaged expression from the target cell’s original RNA profile results in a local perturbation signature, the component of each cell’s transcriptome that specifically reflects its genetic perturbation. Notably, our procedure is capable of characterizing both linear and non-linear perturbation effects, and requires minimal prior knowledge (for example, it does not require a pre-computed list of cell cycle genes). We note that this focuses downstream analyses on changes in expression, rather than cell-state proportions. However, we independently tested for relationships between each perturbation and the resulting fraction of cells in each cell-cycle state, and found no significant effects.

We then repeated principal components analysis and UMAP visualization based on these perturbation signatures, and found that variation in replicate, cell cycle state and activation of cellular stress was substantially mitigated (Figure 2D). As a result, we observed two clear groups of cells expressing a consistent set of gRNAs, including a cluster consisting of cells perturbed for key upstream components of the IFNγ pathway (*IFNGR1, IFNGR2, JAK2, STAT1*), and a second consisting of cells lacking the downstream IFNγ mediator *IRF1*. Cells from the remaining 21 perturbations grouped into a single cluster in this unsupervised analysis. However, cells with a subset of gRNAs (for example, *SMAD4*) were not evenly distributed and showed evidence of substructure (Supplementary Figure 4), suggesting that additional computational improvements may help to clarify their unique molecular perturbations.

**Figure 4.**
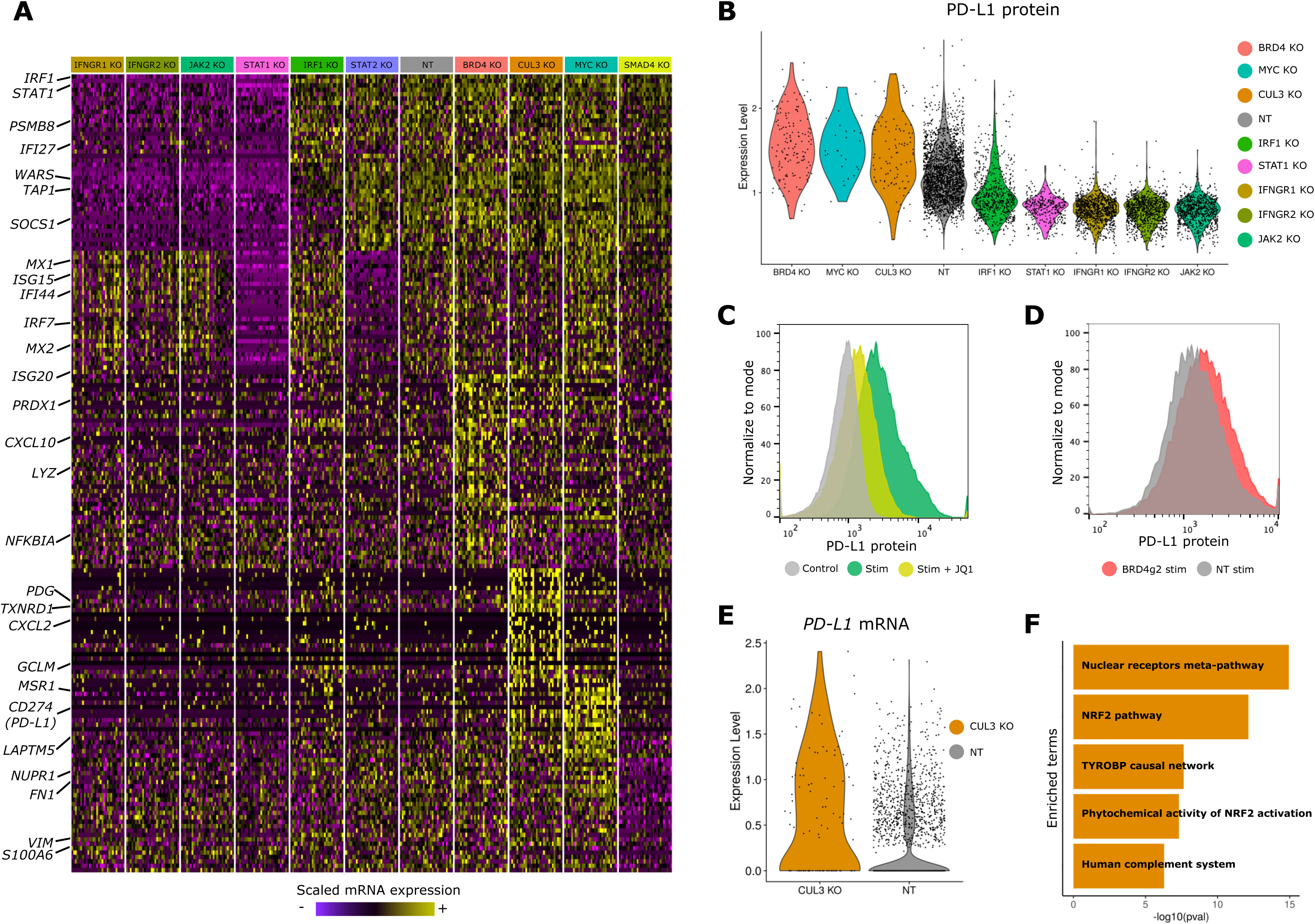
*BRD4* and *CUL3* are negative regulators of PD-L1 expression. **(A)** Single-cell mRNA expression heatmap showing 20 differentially-expressed genes for each mixscape-classified perturbation. For visualization purposes we downsampled our dataset to include 30 cells from each class in the heatmap. **(B)** Violin plots of PD-L1 protein expression for all identified regulators. *BRD4, CUL3* and *MYC* are negative regulators, while the remaining are positive (p-value < 1e^-6^ in all cases). **(C)** Flow cytometry measurements of PD-L1 protein expression across experimental conditions. JQ1 inhibitor treatment (24 hours, 1μM) reduces stimulation-induced PD-L1 expression. **(D)** Flow cytometry measurements of PD-L1 protein expression based on individual gRNA perturbations, validating our ECCITE-seq findings. **(E)** Violin plots showing elevated expression of PD-L1 transcript in CUL3 KO cells, in comparison to non-targeting controls. **(F)** Barplot summarizing gene set enrichment analysis results for 300 genes upregulated in CUL3 KO cells. Analysis was performed using the Human WikiPathways database from the EnrichR package, and reveals a strong enrichment for the *NRF2* pathway.

### A subset of cells ‘escape’ molecular perturbation

The ECCITE-seq data clearly identified the substantial molecular consequences and distinct clustering associated with perturbation of key IFNγ components. For example, IFNGR2g2 cells in the perturbed cluster (circled cells in Figure 2E), exhibited sharp decreases in the expression of hundreds of IFNγ pathway genes, and in *PD-L1* protein levels as well (Figure 2F, G). However, a subset of these cells also appeared to ‘escape’ molecular perturbation. Out of the 1,193 expressing gRNAs targeting *IFNGR2*, 74% were members of the perturbed cluster, but the remaining 26% were indistinguishable from non-targeting controls (Figure 2F, G), demonstrating heterogeneous functional responses among cells expressing the same gRNA.

As has been previously suggested [19,21], cells that ‘escape’ perturbation may not have a deleterious mutation at the Cas9 gene target site. We explored this idea by isolating reads overlapping the IFNGR2g2 gRNA cut site. Since the gene was highly expressed in the ECCITE-seq data, and the gRNA cut site was fortuitously located near the 5’ end, we were able to recover reads for 16,543 cells in the overall dataset (278 of these cells expressed IFNGRg2 gRNA, of which 115 appeared to escape perturbation), and characterized the specific mutations that were introduced. As expected, non-targeted cells did not contain insertions or deletion mutations at the cut site (INDELs), while ‘perturbed’ cells typically exhibited frameshift INDELs (Figure 2H, I). Strikingly, ‘escaping’ cells, when mutated, were primarily characterized by in-frame INDELs, particularly for three or six bases (Figure 2H, I). These results confirm that a substantial fraction of cells escape the introduction of a deleterious mutation, and therefore exhibit no functional consequence of perturbation.

While this phenomenon will also weaken the signal in bulk screens, the ECCITE-seq readout provides us with an opportunity to remove ‘escaping’ cells from the analysis. Due to the limited depth of scRNA-seq based readouts (alongside the ability to profile mutations outside the transcript end), we cannot directly measure the mutational profile of each cell in the vast majority of cases. However, inspired by previous pioneering work [19,21,26], we reasoned that we could use the cell’s transcriptome as a phenotypic readout of the presence or absence of a deleterious mutation, and developed a strategy to systematically identify and remove ‘escaping’ cells.

### *Mixscape* robustly classifies ‘non-perturbed’ cells

Our analytical solution to identify ‘escaping’ cells is inspired by a classification tool known as Mixture Discriminant Analysis (MDA). MDA assumes that individual samples fall into different groups, but that each group is a mixture of *n* different subclasses [27]. This assumption is valid for our ECCITE-seq data, where individual cells can be divided into groups dependent on their expressed gRNA, but each group can represent a mixture of ‘perturbed’ and ‘escaping’ (or non-perturbed) subclasses. MDA fits Gaussian mixture models for data points in each group, enabling the assignment of subclass identity.

We therefore modeled our ECCITE-seq transcriptomic data using a mixture of Gaussians, but placed two constraints on the method. First, we set *n*=1 for the ‘control’ group, and *n*=2 for all other gRNA-defined groups. Second, based on our previous observations (Figure 2E-G), we assumed that the ‘escaping’ cells exhibit a perturbation signature that is similar to ‘control’ cells. When fitting Gaussian mixture models, we therefore constrained the parameters for one of the mixture components to mirror the ‘control’ cells. We refer to the resulting procedure as *mixscape*. For each targeted cell, *mixscape* considers a cell’s perturbation signature (calculated as previously described), and assigns it to a ‘perturbed’ or ‘escaping’ subclass (Figure 3A).

We validated the *mixscape* predictions on IFNGR2 cells (74.6% classified as perturbed (‘KO’), 25.4% classified as non-perturbed (NP), i.e. an 74.6% perturbation rate), by confirming that only cells predicted as KO exhibited reductions in IFNγ target expression and PD-L1 surface protein levels. We observed similar results for additional interferon-regulators, including *IFNGR1, IFNGR2, JAK2, STAT1*, and *IRF1* (Figure 3B). Interestingly, *mixscape* predicted substantial variation in the perturbation rate of four independent *IRF1* gRNAs, ranging from 39% to 92% (Figure 3C, black boxes). To independently measure the efficacy of each guide, we used flow cytometry to assess its effect on PD-L1 protein expression (Figure 3D, E). These measurements were concordant with *mixscape* predictions, further validating our approach.

We note that in cases where functional removal of a gene fails to result in a detectable transcriptomic shift, *mixscape* will also mark a cell as non-perturbed, even if a frameshift mutation was introduced (Supplementary Figure 6A-C). Indeed, for 15 genes, *mixscape* predicted a 0% perturbation rate. In each of these cases, we also found no differentially expressed genes when comparing cells targeted by these gRNA to non-targeted controls. Furthermore, when we attempted to classify cells expressing a NT gRNA as a negative control, *mixscape* correctly predicted a 0% perturbation rate. Importantly, these results demonstrated that *mixscape* does not overfit the data and only predicts cells to be in the ‘perturbed’ class when there is a detectable change in their molecular state.

A full description of *mixscape* is presented in the Supplementary Methods, alongside comparative benchmarking with MIMOSCA [19] and MUSIC [26] (Supplementary Figures 7 A-D and 8 A-D). We used both positive controls (cells targeted with the IFNGRg2 guide) and negative controls (cells targeted with a NT gRNA) to evaluate performance, and found that *mixscape* was the only method capable of sensitively identifying perturbed cells without overfitting (Supplementary Figures 7A and 8A). We have implemented *mixscape* as part of Seurat, our open-source R toolkit for single-cell analysis [28], and include an introductory vignette (Supplementary Note 1) demonstrating how to run the software on our ECCITE-seq dataset.

For 11 genes, *mixscape* did predict the presence of perturbations, with a perturbation rate varying from 23% to 83%. This variation could reflect differences in the targeting efficiency of individual gRNAs, the strength of perturbation for each individual gene, or differences in the dosage requirement (heterozygous vs homozygous KO) for each putative regulator. We also note that our observed perturbation rate could be skewed for perturbations that result in cell death, as these could selectively deplete KO cells. Regardless, these analyses highlight the importance of characterizing the extensive heterogeneity within cells that receive the same sgRNA. In downstream analyses, we chose to remove cells that were predicted to escape perturbation, as including these cells will substantially dampen the biological effects associated with gene knockout.

To visualize the remaining 11 classes we applied Linear Discriminant Analysis (LDA). LDA aims to identify discriminant functions that maximally differentiate the *mixscape*-derived classes (Supplementary Methods). We then used these discriminant functions as input to generate a two-dimensional UMAP for visualization (Figure 3F). We found that the resulting UMAP effectively separated the different perturbations, with the exception of a negative control (Supplementary Methods), while maintaining local proximity for similar perturbations (i.e. cells targeted with gRNA against *IFNGR1* and *IFNGR2* are adjacent in the embedding). Using LDA as an initial step improved separation in all cases except for the negative control (Supplementary Figure 9), suggesting that combining LDA with UMAP is an effective approach for the visualization of pooled single-cell sequencing screens.

### CUL3 and BRD4 are negative regulators of PD-L1 expression

These analyses suggest that after removing non-perturbed cells, each genetic knockout induces a specific molecular response. Indeed, when performing differential expression compared to control cells, we observed striking differences in gene expression that defined each molecular perturbation (Figure 4A). Of particular interest, we observed that perturbation of eight genes also resulted in a shift of PD-L1 protein levels in our ECCITE-seq data (Figure 4B). We identified five positive regulators (*PD-L1* downregulation upon perturbation) and three negative regulators, a subset of which had been previously validated [9,11,13,16,17,29]. For example, in addition to the core components of the IFNγ pathway, we verified that perturbation of BHLH transcription factor MYC [12] and the ubiquitin ligase CUL3 [15] both increase PD-L1 surface protein levels, consistent with previous reports. These results demonstrate the potential for ECCITE-seq data to robustly and accurately characterize multiplexed perturbations. Importantly, perturbation of these eight genes did not result in appreciable shifts in CD86 and PDL2 protein expression (Supplementary Figure 10A-B) suggesting that these regulatory effects are specific to PD-L1.

To our surprise, we observed that perturbation of the bromodomain-containing protein *BRD4* resulted in a upregulation of PD-L1 protein levels, indicating that BRD4 acts as a negative regulator. Previous studies have utilized the bromodomain inhibitor JQ1, an alternative to *BRD4* genetic perturbation, to suggest that *BRD4* is in fact a positive regulator of *PD-L1* [13,29]. To help reconcile these differences, we treated our stimulated cells with JQ1 and observed a reduction in PD-L1 expression (Figure 4C). However, we validated that CRISPR-mediated genetic perturbation of BRD4 leads to an up-regulation of PD-L1 expression using flow cytometry (Figure 4D), confirming the ECCITE-seq result. These results indicate that BRD4 is a negative regulator of PD-L1 expression, and that the JQ1 inhibitor may interact with additional proteins in order to achieve PD-L1 reduction.

We also observed that *CUL3* and *BRD4* perturbation resulted in similar levels of PD-L1 protein upregulation (Figure 4B). To our surprise, while the ubiquitin ligase complex CUL3-SPOP has been shown to post-transcriptionally regulate PD-L1 protein levels [15], we also detected a 1.6-fold (p < 10^−11^) upregulation of *PD-L1* mRNA levels (Figure 4E). We observed both protein and mRNA up-regulation only in cells predicted to be perturbed by *mixscape*. Our results suggest that in addition to its known role in regulating PD-L1 protein stability via direct ubiquitination, *CUL3* perturbation also modulates PD-L1 mRNA levels.

To gain further insight into the effects of *CUL3* perturbation, we identified differentially expressed genes (DE) between *CUL3*-perturbed and control cells, and intersected these genes with members of previously identified transcriptional pathways (Supplementary Figures 11A, B). We observed no overlap with canonical IFNγ signaling targets, suggesting that CUL3-mediated transcriptional regulation of *PD-L1* is mediated through an IFNγ-independent pathway. Instead, we observed a striking enrichment (p < 10^−14^) for target genes of the Nuclear factor erythroid-2 factor 2 (*NRF2*) signaling pathway (Figure 4F).

### *CUL3* indirectly regulates *PD-L1* at the transcriptional level through *NRF2*

The *NRF2* pathway is activated during oxidative stress, and induces the expression of many antioxidant genes to prevent cellular damage and death [30]. *NRF2* has been shown to directly bind to the *PD-L1* promoter and activate transcription under ultraviolet-induced stress [14], and NRF2 protein stability is directly regulated by the CUL3-KEAP1 ubiquitin ligase complex [31]. Taken together with these findings, our data suggest that CUL3 may have two distinct mechanisms for regulating PD-L1 protein expression. First, as previously described [15], perturbation of the CUL3-SPOP complex interferes with the ubiquitination of PD-L1, directly enhancing its stability and protein expression level. Second, our data indicate that perturbing the CUL3-KEAP1 complex interferes with the ubiquitination of NRF2, boosting pathway activation and *PD-L1* transcript expression (Figure 5A).

**Figure 5.**
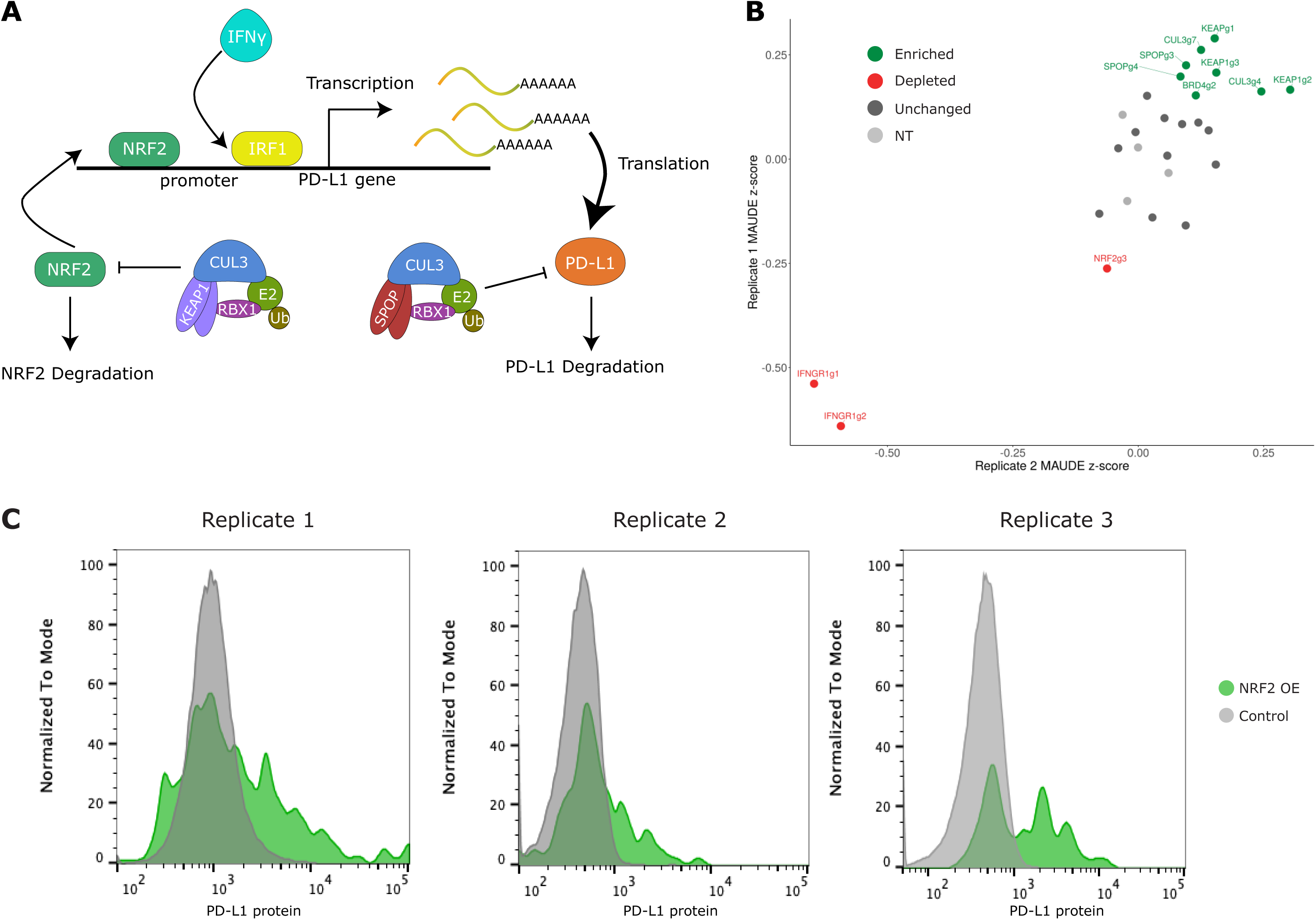
CUL3-KEAP1 complex indirectly regulates *PD-L1* transcript levels by regulating NRF2 protein stability. **(A)** Schematic representation describing two complementary modes of *CUL3*-mediated *PD-L1* regulation. The CUL3-SPOP complex directly regulates PD-L1 protein stability through ubiquitination. The CUL3-KEAP1 complex regulates NRF2 protein stability, indirectly modulating NRF2-mediated *PD-L1* transcription. **(B)** Validation pooled CRISPR screen results (2 biological replicates) targeting *KEAP1, SPOP, CUL3, BRD4, IFNGR1* and *NRF2* (including 4 non-targeting gRNAs). gRNAs targeting *KEAP1, SPOP, CUL3* and *BRD4* (green) were enriched in cells expressing high levels of PD-L1 protein while *NRF2* and *IFNGR1* gRNAs were depleted (red). **(C)** Flow cytometry measurements of PD-L1 protein expression in *NRF2*-overexpressing (green) or control (grey) THP-1 cells. Overexpression of *NRF2* results in upregulation of PD-L1 protein when compared to control cells. Three independent replicates are shown.

In order to validate that CUL3 acts as an indirect regulator of *PD-L1* mRNA levels, we performed a focused validation screen by infecting cells with 27 gRNAs targeting 6 genes (Supplementary Table 1). We used flow cytometry to isolate bins of PD-L1 high (PD-L1^hi^) and low expressing (PD-L1^lo^) cells after stimulation (Supplementary Figure 12), sequenced the gRNA locus for each bin, and compared the gRNA representation. gRNAs against genes that were predicted to be negative regulators of PD-L1, including *CUL*3, and *KEAP1* were consistently overrepresented in PD-L1^hi^ cells in two biological replicates (Figure 5B), while we observed the converse for predicted positive regulators (*NRF2* and *IFNGR1*).

As an independent validation, we found that direct overexpression of *NRF2* in THP-1 cells resulted in an up-regulation of PD-L1 protein by flow cytometry (Figure 5C). Taken together, our data demonstrate that by modifying the activity of the *NRF2* pathway, the CUL3-KEAP1 complex is an indirect regulator of PD-L1, and highlight the potential for ECCITE-seq to disentangle complex regulatory pathways via simultaneous characterization of both RNA and protein modalities.

## DISCUSSION

In this study, we coupled pooled CRISPR screens to a multi-modal single-cell sequencing readout in order to investigate the regulation of IC proteins, such as PD-L1. We leveraged our dataset to characterize the transcriptional and post-transcriptional effects of 111 independent perturbations. To assist in this process, we developed unsupervised computational methods to control for confounding sources of variation that can mask perturbation signals in ECCITE-seq datasets. Our analyses identified numerous regulators of PD-L1 expression, and in particular, two negative regulators (BRD4 and CUL3) which we validated using complementary approaches.

The multi-modal nature of ECCITE-seq data enabled us to move beyond the identification of regulators towards a more in-depth molecular characterization. For example, we found that CUL3-KEAP1 can act as an indirect regulator of *PD-L1* mRNA levels, in addition to the previously identified role for CUL3-SPOP in directly regulating PD-L1 protein stability. These findings are intriguing in light of recent reports that *KEAP1* is often mutated in lung cancer, and mutations in the *NRF2/KEAP1* have been associated with treatment resistance [23,32]. Future studies may benefit from exploring possible links between these mutations and the expression of IC molecules.

Our datasets also highlight that cells which are targeted with the same sgRNA are inherently heterogeneous. First, we demonstrated that the calculation of a ‘local’ perturbation signature can remove confounding sources of variation from downstream analyses, even when these sources are unknown. Second, we introduce *mixscape*, inspired by mixture discriminant analysis and building on previous pioneering methods [19,21,26]. *Mixscape* robustly filters cells that do not exhibit transcriptomic evidence of perturbation, and substantially increases the signal/noise ratio in downstream analyses. The ability to computationally leverage the heterogeneity within targeted cells is a distinct advantage of coupling genetic screens to a single-cell sequencing readout. Importantly, alternative genetic perturbations such as CRISPR interference and CRISPR activation may reduce this heterogeneity, though confounding sources of variation and ‘escaping’ cells are likely to characterize these technologies as well.

One limitation of *mixscape* is the reliance on detecting a shift in gene expression in order to classify cells. In particular, perturbations that modify alternative phenotypes, such as epigenetic state, protein levels, or functional responses, but exhibit no evidence of transcriptomic change will be classified as ‘no detected perturbation / non-perturbed (NP)’. In this manuscript, we inferred perturbation status using the transcriptome, and validated our calls using surface protein levels from ECCITE-seq. However, integrative multi-modal approaches [33] could enable joint analysis of the transcriptome and protein levels when filtering NP cells, and represent a promising future extension of our method.

Lastly, we note that *mixscape*’s binary classification of targeted cells likely represents an oversimplification that can be improved with additional experimental data from large-scale future experiments. Genetic perturbation with CRISPR/Cas9 introduces a diverse set at the cut site. As datasets increase in size, we envision sufficient scale to characterize how each precise mutation has a unique (though potentially subtle) effect on a cell’s molecular phenotype. Moreover, rapid molecular advances continue to enable the simultaneous measurement of additional cellular components, such as chromatin state and gene expression [34–37]. Together, these data will enable systematic perturbation of gene structure and dosage, alongside detailed characterization of multiple molecular modalities.

## AUTHOR CONTRIBUTIONS

EP, EM, PS, and RS conceived the research. EP, EM, SF, BB, WMM, HHW, BZY, and PS performed experimental work. EP, AWB, and RS performed computational analyses. All authors participated in interpretation and in writing the manuscript.

## ACKNOWLEDGEMENTS

We acknowledge Ross Levine, Thales Papagiannakopoulos, and members of the Satija and Technology Innovation Labs at NYGC for general discussion, Patrick Roelli for assistance with pre-processing, and Neil Bapodra and Neville Sanjana for advice on vector and library design. This work was supported by the Chan Zuckerberg Initiative (EOSS-0000000082 to RS, HCA-A-1704-01895 to PS and RS), and the National Institutes of Health (DP2HG009623-01 to RS, RM1HG011014-01 to PS and RS, R21HG009748-03 to PS).

## DATA AND CODE AVAILABILITY

Raw and processed sequencing data is available through the Gene Expression Omnibus (GEO accession number: GSE153056). Processed data is also available through SeuratData (https://github.com/satijalab/seurat-data) to facilitate access with a single command: InstallData(ds = “thp1.eccite”). The code for *mixscape* is freely available as open source software as part of the Seurat package for single-cell analysis (www.github.com/satijalab/seurat/tree/mixscape). A vignette demonstrating the application of *mixscape* to this dataset is available as a supplementary note (Supplementary Note 1), as well as an online resource (https://satijalab.org/seurat/vignettes/mixscape_vignette.html)

## SUPPLEMENTARY METHODS

### Cell culture and Maintenance

THP-1 cell line was obtained from ATCC (TIB-202) and was grown at 37C in RPMI medium supplemented with 10% FBS. To induce the expression of various immune checkpoint proteins cells were treated with Decitabine (Sigma-Aldrich A3656, 0.25μM) for three days, TGFβ1 for two days (Thermo Fisher Scientific PHG9204, 2.5ng/ml) and IFNγ for one day (R&D systems 284-IF-100, 10ng/ml). HEK293FT human embryonic kidney (#R70007) cells were grown in DMEM medium supplemented with 10% FBS (D10). The D10 medium for HEK293FT cells was additionally supplemented with 6mM L-glutamine (Thermo Fisher Scientific, #25030081), 1mM Sodium Pyruvate (Thermo Fisher Scientific, #11360070) and 0.1mM MEM Non Essential Amino Acids (Thermo Fisher Scientific, #11140050). TrypLE (Thermo Fisher Scientific, #12604039) was used to lift HEK293FT cells from plates during passaging. All cells were passaged every two to three days and low passage cells were used for all experiments (p3-p12).

### Flow Cytometry

After treatment, cells were centrifuged at 300g for five minutes and resuspended in 100μl of MACS buffer (1X PBS, 0.5% BSA, 2mM EDTA). 5μl of FcX blocking reagent was added and cells were placed on ice for 10 minutes. Next, antibodies were added directly into the mix and cells were kept on ice for another 30 minutes. Prior to flow cytometry (FACS), cells were passed through a 40μm cell strainer (VWR, #10032-802) to remove any cell clumps. The following FACS antibodies were used in these experiments at concentrations recommended by the manufacturer: PD-L1 (BD Biosciences, #558017), PD-L2 (BioLegend, #329606), CD86 (BioLegend, #305412). Compensation beads were used to overcome signal overlap between fluorophores (BD Biosciences, #552843). To check and remove any dead or apoptotic cells DAPI (Sigma Aldrich, #D9542-5MG) was added to the staining mix at a concentration of (0.4μg/1mL). All FACS measurements were performed using the SONY SH800 cell sorter. FACS analyses and plots were made using the FlowJo™Software [38].

### CITE-seq experiment

THP-1 cells were stimulated as described above or left unstimulated. At the end of the stimulation, cells were collected by centrifugation at 300g for 5 minutes. Cells were resuspended in 100μl of staining buffer containing 5μl of FcX blocking reagent and were placed on ice for 10 minutes. Next, 100μl of staining buffer containing CITE-seq antibodies (0.5μg/antibody/sample) was added to the cells. The cells were placed in the 4C fridge for 30 minutes to allow for antibodies to bind to their target protein. For the CITE-seq experiment antibodies were conjugated in-house following the hyper Oligo-antibody conjugation protocol as detailed here (https://cite-seq.com/protocol/). To keep track of the experimental condition (stimulated vs unstimulated) and be able to detect and remove cell doublets, cells were aliquoted into three tubes containing a uniquely barcoded hashing antibody. Cells were placed in the fridge for an additional 20 minutes. After staining was complete, all samples were washed three times with 1ml staining buffer to remove all the excess unbound antibodies. Next, cells were resuspended in 200-300μl of 1X PBS and counted using the Countess II Automated cell counter system. Immediately before loading to the 10x Genomics instrument, cells from all experimental conditions were pooled at the appropriate concentration (recovery of 10,000 cells per lane).

### CITE-seq data library construction, sequencing and data analyses

We ran 1 lane of 10x Genomics 5’ (Chromium Single Cell Immune Profiling Solution v1.0, #1000014, #1000020, #1000151) aiming for 20,000 cell recovery per lane. Prior to the run, cell viability was determined and cell numbers were estimated as previously described. To increase the number of cells assayed we hashed them following the cell hashing protocol [25]. mRNA, hashtags (Hashtag-derived oligos, HTOs) and protein (Antibody-derived oligos, ADTs) libraries were constructed by following 10x genomics and CITE-seq protocols. All libraries were sequenced together on a Novaseq run. Sequencing reads coming from the mRNA library were mapped to the *hg19* reference genome using the *Cellranger* Software (V2.1.0). To generate count matrices for HTO and ADT libraries, the *CITE-seq-count* package was used (https://github.com/Hoohm/CITE-seq-Count). Count matrices were then used as input into the *Seurat* R package [28,39] to perform all downstream analyses.

Cells with low quality metrics, high mitochondrial gene content (> 10%) and low number of genes detected (< 500) were removed. RNA counts were log-normalized using the standard Seurat workflow. ADT and HTO counts were normalized using the centered log ratio transformation approach, with a *margin = 2* (to normalize across cells instead of across features). To identity cell doublets and assign experimental conditions to cells, we used the *HTODemux* function. We performed PCA on the protein measurements, observing a continuum in the level of PD-L1 up-regulation, and selected the top 200 genes whose expression correlated with this continuum. These genes are shown in Figure 1D, where cells in both the protein and RNA heatmaps are ordered based on their PC1 embedding values.

### CITE03 plasmid construction

To increase sgRNA targeting efficiency we switched the sgRNA scaffold on the CROP-seq plasmid (addgene, #86708) with the optimized sgRNA scaffold as described in [40]. Moreover, we replaced the puromycin resistance gene on the CROP-seq plasmid with a blasticidin resistance gene fused to eGFP amplified from the pFUGW-EFS-V5-EGFP-2A-Bla-WPRE plasmid (addgene, #71215). Finally, we removed Cas9 protein to decrease the size of our plasmid and achieve higher viral titer.

### Inducible Cas9 THP-1 cell line

The THP-1-Cas9 inducible cell line was made by lentiviral transduction using the pCW-Cas9-puro plasmid (addgene, #50661). Single cells were sorted into 96-well plates three days after puromycin selection to obtain single cell clones. Single cell colonies were expanded for four weeks before assessing Cas9 expression. Protein lysates were obtained from ten clones before and after 24hrs of doxycycline treatment (1μg/ml, Sigma-Aldrich D9891) to check Cas9 expression by westernblot. Briefly, cells were washed 2 times with 1mL of ice-cold 1X PBS and resuspended in RIPA lysis buffer (Amresco, N653) supplemented with a protease inhibitor cocktail (Bimake, B14001). Cas9 expression was verified by western blot using GAPDH antibody as loading control (Cell signaling Technology, 2118S) and Flag antibody (Cell signaling Technology, 14793S) to detect Cas9 protein. Protein bands were visualized using fluorescently labeled secondary antibodies (LI-COR, #925-32212 and #925-68073) and the Odyssey Imaging System. One of the clones with the highest Cas9 expression was selected and used for all downstream experiments. To minimize leakiness of our doxycycline inducible Cas9 system, a TET-free FBS (VWR, 97065-310) was used to grow these cells.

### gRNA design, virus production and Cas9 dynamics

*Guides* webtool (http://guides.sanjanalab.org/#/) was used to predict gRNAs with high targeting efficiency and low off-target effects [41]. 3-4 guides per gene were selected together with 10 guides predicted to have no sequence similarity with the human genome (non-targeting controls). Guide oligos were synthesized individually using IDT. Oligos were cloned into the CITE03 vector as previously described [42]. Low passage HEK293FT cells were transfected with MD2.G (addgene #12259), PAX2 (addgene #35002) and the CITE03 plasmids carrying gRNAs using Lipofectamine 2000 (Thermo Fisher Scientific, #11668030). Media was replaced with DMEM + 10% FBS + 1% BSA (NEB, B9000S), 6 hours post-transfection. Viral supernatants were harvested 48-72 hours post transfection by centrifugation (ten minutes, 3000 rpm, 4C) and stored in a -80C freezer until used. To estimate the concentration of the virus, cells were infected with increasing amounts of virus and three days post antibiotic selection, the percentage of dead and live cells was calculated. In all experiments, cells were infected at low multiplicity of infection (MOI) to achieve one gRNA insertion per cell.

To estimate how many days after Cas9 induction we have saturation of CRISPR-induced insertions and deletions (INDELs), we ran single gRNA experiments targeting PD-L1 protein. Cas9 was induced with the addition of doxycycline (1μg/mL) for one, three, five and seven days and we used TIDE [43] and Surveyor assays (IDT, #706020) to estimate the percentage of cells with INDELs. As an independent method, we also used flow cytometry to check PD-L1 expression and quantify the percentage of knockout cells (KO). We found that after five days of Cas9 induction the percentage of cells with INDELs stops increasing and we have achieved the highest percentage of cells with low PD-L1 protein expression. Based on these observations, we decided to treat cells with 1μg/mL of doxycycline for five days prior to running the ECCITE-seq experiments.

### ECCITE-seq pilot experiment

We ran an initial pilot experiment to validate our ability to accurately recover gRNA and plan experimental design. We generated single gRNA cell lines for 20 gRNA, including PD-L1, IFNGR1, and non-targeting controls, and performed individual infections. Next, we stimulated cells as previously described. We hashed each cell line separately [25] prior to running our ECCITE-seq experiment. This experimental set up enabled us to have two independent methods for encoding the perturbation received by each cell. Libraries were sequenced on a NextSeq500. mRNA libraries were quantified using Cell Ranger (2.1.1; hg19 reference), and normalized using standard log-normalization in Seurat. HTO and ADT libraries were processed with *CITE-seq-count* (https://github.com/Hoohm/CITE-seq-Count), and normalized using the centered log-ratio (CLR, across cells). Cells with high mitochondrial gene content (> 8%) were removed. RNA counts were log-normalized using the standard Seurat workflow. ADT, HTO and GDO counts were normalized using the centered log ratio transformation approach, with a *margin = 2* (to normalize across cells instead of across features).

We demultiplexed the cell hashing data using the *MULTIseqDemux* function adopted from [44], and removed all classified doublets. We assigned gRNA identity using HTODemux in Seurat. To assess the accuracy of gRNA classification, we examined each cell with an identified gRNA, and compared its classification to its HTO-derived label. We observed an overall concordance of 99.4%. Concordant cells were used for plotting PD-L1 expression in Figure 1E.

### ECCITE-seq experimental setup

THP-1 Cas9-inducible cells were transduced with virus containing 111 guides at low MOI to obtain cells with 1 gRNA. 24 hours post-transduction cells were centrifuged and resuspended in new media containing blasticidin (15μg/mL) to select for successfully transduced cells. Three days after antibiotic selection, media was exchanged with fresh R10 containing blasticidin (15μg/mL) and doxycycline (1μg/mL) to induce Cas9 expression and INDEL formation. After five days of doxycycline treatment, cells were stimulated with DAC, IFNγ and TGFβ1 for an additional three days or left unstimulated prior to running the 10x Genomics experiment (Supplementary Figure 2A). The final pool of cells loaded onto the 10x Genomics chip contained 10% of unstimulated cells and 90% of stimulated cells coming from four biological replicates.

### Single cell ECCITE-seq library construction and sequencing

For the ECCITE-seq experiment, we run eight lanes of 10x Genomics 5’ (Chromium Single Cell Immune Profiling Solution v1.0, #1000014, #1000020, #1000151) aiming for 10,000 cell recovery per lane. Prior to the run, cell viability was determined and cell numbers were estimated as previously described. To keep track of each biological replicate identity, samples were hashed following the cell hashing protocol [45]. mRNA, hashtags (Hashtag-derived oligos, HTOs), protein (Antibody-derived oligos, ADTs) and gRNA (Guide-derived oligos, GDOs) libraries were constructed by following 10x genomics and ECCITE-seq protocols. All libraries were sequenced together on two lanes of a NovaSeq run. Sequencing reads coming from the mRNA library were mapped to the *hg19* reference genome using the *Cellranger* Software (V2.1.1). To generate count matrices for HTO, ADT and GDO libraries, the *CITE-seq-count* package was used (https://github.com/Hoohm/CITE-seq-Count). Count matrices were then used as input into the *Seurat* R package [28,39] to perform all downstream analyses.

### ECCITE-seq data pre-processing in *Seurat*

Cells with low quality metrics, high mitochondrial gene content (> 10%) and low number of genes detected (< 100) were removed. RNA counts were log-normalized using the standard Seurat workflow. ADT, HTO and GDO counts were normalized using the centered log-ratio transformation approach, with *margin = 2* (normalizing across cells). To identity cell doublets and assign experimental conditions to cells, we used the *MULTIseqDemux* function adopted from [44]. MULTIseqDemux-defined cell doublets and negatives were removed from any downstream analyses. To assign a gRNA identity to each cell, we looked at the GDO counts. If a cell had less than five counts for all gRNA sequences we classified it as negative. For all other cells, we found the gRNA with the highest number of counts and assigned it to that cell. Cells that had high counts for more than one gRNA were classified as doublets.

We checked the gRNA representation across all four biological replicates included in this experiment by calculating the percentage of cells that belonged to each gRNA class within each biological replicate (Supplementary Figure 2B). We removed replicate #4 (both stimulated and unstimulated cells) as it had a skewed gRNA representation, likely due to long term cell culture. We also removed cells in target gene classes where less than 10 total cells were detected, even after pooling across gRNA and replicates.

### RNA-based clustering of single cells

To visualize cells based on an unsupervised transcriptomic analysis (Figure 2A), we first ran PCA using 2000 variable genes. The first 40 components were used as input for UMAP visualization in two-dimensions [46]. We calculated cell-cycle scores using the CellCycleScoring function in Seurat v3.1 with default parameters.

### Calculating perturbation signatures for single cells

Let *X* = {*x*_1_,…, *x*_*N*_} represent a normalized single-cell dataset, with *N* cells.

For each cell *x*_*i*_, we perform the following procedure:

1. Identify *Y*_*i*_, a subset of *X* consisting of cells that receive a ‘non-targeting’ gRNA, and were present in the same biological replicate *r* as *x*_*i*_
2. Identify the set {*y*_*i,1*_,*…,y*_*i,k*_}of nearest neighbors to *x*_*i*_, based on the top 40 principal components described above. We set the hyperparameter *k* = 20 by default, and identified neighbors using the Randomized Approximate Nearest Neighbors (RANN) algorithm [47].
3. Compute the average expression profile of this local neighborhood 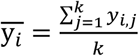
4. Compute the ‘local’ perturbation signature 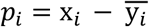

This calculation is implemented in the CalcPerturbScore function in Seurat.

### Clustering single cells based on their perturbation signature

Perturbation signatures were centered but not scaled using the ScaleData() function. We ran PCA using the perturbation signatures of the top 2000 most variable genes defined using the RNA assay. The first 40 components were used as input for UMAP visualization in two-dimensions [46].

### Estimating % of INDELs from scRNA-seq reads

We used Sinto (https://timoast.github.io/sinto/basic_usage.html) to extract all sequencing reads that belonged to the perturbed and non-perturbed IFNGR2g2 cells as well as the non-targeting control cells from the cellranger *possorted genome bam* files. Bam files from all 10x Genomics lanes were merged to three final bam files, one for each group (Non-targeting, knockout and non-perturbed). Samtools [48] was used to create the index file used for visualization into IGV tools Software [49]. To quantify the percentage of INDELs at the expected gRNA cut site, we used GenomicRanges, GenomicFeatures, GenomicAlignments, Rsamtools and bedr R packages. First, a bed file was constructed to specify the gRNA cut site. Next, we removed any reads that didn’t overlap our cut site. To ensure accurate INDEL quantification, we only assessed reades that extended enough into the 3’ end of the gRNA sequence. We relied on the read cigar string information to quantify the number of reads with frameshift or inframe mutations by looking at the number of bases inserted/deleted (three or multiple of three = inframe, any other as frameshift). To calculate the percentage of inframe and frameshift deletions we divided each class by the total number of reads post filtering.

### Mixture-model based classification of KO and NP cells

The objective of this procedure is to identify cells that received a targeting guide but exhibited no detectable transcriptomic evidence of perturbation. We perform the following procedure independently, for each targeted gene *g*.

1. We perform differential expression testing between all cells that receive gRNA targeting gene *g*, and all cells that receive a NT gRNA. The gene set *DEG* represents the set of genes that pass a Bonferroni-adjusted p-value threshold of 0.05. If *DEG* consists of fewer than five genes, we stop the procedure, and label all cells as non-perturbed.
2. *Let* 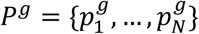, represent a set of single-cell perturbation signatures for *N* cells, each of which receives gRNA targeting gene *g*. Similarly, let 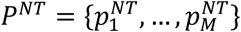, represent a set of single-cell perturbation signatures for all *M* cells that receive a non-targeting gRNA.
3. For each cell, the perturbation signature is a vector, with length equal to the size of the *DEG* gene set. We project this into a single dimension, representing a perturbation score *s* for each cell. We find that reducing the dimensionality of these data substantially improves the robustness to overfitting in downstream analyses. To calculate the score, we first calculate a vector representing the difference in the average perturbation signature of targeted and non-targeted cells. We then project each cell’s perturbation signature onto this vector. Specifically:

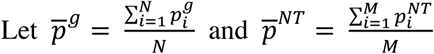

Then the perturbation score *s* for cell *i* is defined by:

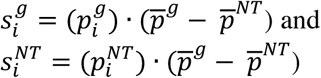
4. We model the perturbation scores of non-targeting cells with a Gaussian distribution:

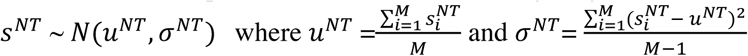
5. We model the perturbation scores of targeted cells using a mixture of two Gaussian distributions. One mode represents cells that resemble NT cells due to a lack of a detectable perturbation, and therefore is parameterized by the previously measured *U*^*NT*^ and *σ*^*NT*^

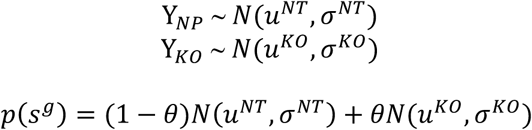

This requires estimating three parameters: the mean and standard deviation rate for the perturbation score of KO cells (*u*^*KO*^, *σ ^KO^*) and the mixing rate (or ‘perturbation rate’) *θ*. We learn these parameters using the function *normalmixEM* from the *mixtools* package.
6. We calculate the probability that each cell *i* was successfully perturbed by a gRNA targeting gene *g* :

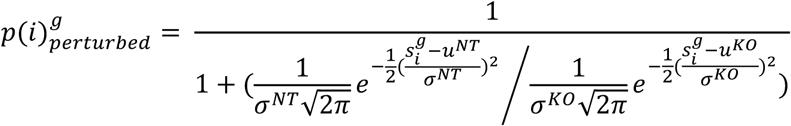
7. All targeted cells with a perturbation probability > 0.5 are classified as KO cells, while the remainder of cells are classified as NT cells.
8. We repeat steps 1-7 until the classifications converge. In this manuscript, all analyses converged within 5 iterations.

At the conclusion of this procedure, each cell is assigned one of three identities:

- If the cell received a NT gRNA, it retains its assignment as non-targeting (NT)
- If the cell received a targeting gRNA, and is classified in step 8 as NT, it is assigned a non-perturbed (NP) label
- If the cell received a targeting gRNA, and is classified in step 8 as KO, it receives a perturbed/knock-out (KO) label.

In addition to returning a KO or NP label, *mixscape* returns a perturbation probability (as defined in step 7) for each targeted cell.

This calculation is implemented in the RunMixscape function in Seurat

### Benchmarking *mixscape* against MIMOSCA and MUSIC

MIMOSCA [19] and MUSIC [26] provide alternative computational frameworks for identifying non-perturbed cells in single cell pooled CRISPR screens. We ran MIMOSCA using the model-fitting procedure with default parameters, as specified in the ‘Computational Workflow’ section of the Github repository README (https://github.com/asncd/MIMOSCA). MIMOSCA requires a gene expression matrix and a file with all target gene classifications. For consistency, our gene expression matrix consisted of all genes used to build our *mixscape* classification model. To run MIMOSCA we used default parameters, which represented optimized values as described in the Perturb-seq publication (sklearn.linear_model.ElasticNet(l1_ratio = 0.5, alpha = 0.0005, max_iter = 10000).

Similarly to MIMOSCA, MUSIC requires the gene expression matrix and a file with all target gene classifications. For consistency, our gene expression matrix consisted of all genes used to build our *mixscape* classification model. MUSIC performs QC to remove low quality cells, and runs SAVER [50] on the gene expression matrix to impute mRNA expression values. The newly imputed matrix, together with the provided classifications, are used to classify cells. We ran MUSIC following the illustratrated example in the Github repository README (https://github.com/bm2-lab/MUSIC, with default parameters.

For benchmarking analyses, prior to running the three methods, we randomly sampled 1,000 cells expressing NT gRNA and re-labeled them as a new targeted gene class, representing a negative control (NEG CTRL). These cells should all be classified as NP (Supplementary Figures 7,8).

### Linear Discriminant Analysis-based dimensionality reduction

After removing non-perturbed cells, we apply Linear Discriminant Analysis (LDA), followed by UMAP [46], to visualize the remaining cells in two dimensions. We apply LDA as an alternative linear reduction technique to PCA. While PCA aims to identify a low-dimensional subspace that maximally retains variation in a dataset, LDA aims to identify a low-dimensional subspace that maximally discriminates different groups of the data. In our case, the input to LDA is a single-cell data matrix and a set of group labels (the mixscape-derived classes).

In principle, we can use normalized gene expression as an input data matrix to LDA. However, this approach can lead to overfitting, as the total number of genes may be of a similar magnitude to the total number of cells. We therefore aimed to first reduce the dimensionality of our data in an unsupervised way, while retaining the sources of variation that distinguished each perturbation. We performed the following procedure for each targeted gene *g*:

1. From the previously computed set of perturbation signatures *P*, we extract all cells that are labeled by mixscape as KO for gene *g*, along with all non-targeted cells.
2. We perform unsupervised PCA. As input features to PCA, we use the gene set DEG, as previously calculated during mixscape classification.
3. We project this subspace onto all cells in the dataset.
4. We retain the first 10 projected components for all cells, and expect that this subspace will retain differences between KO and NT cells.

At the conclusion of this procedure, all retained components are used as input to linear discriminant analysis using the lda function in the MASS R package [51]. This procedure is implemented in the MixscapeLDA function in Seurat.

The results from this function are used as input for 2D visualization with UMAP (Figure 3F). We found that this procedure substantially improved the visualization and interpretability of ECCITE-seq data. We observed that cells characterized by different perturbations separated visually in the 2D embedding, but retained their global structure (for example, STAT1, JAK2, IFNGR1 and IFNGR2 are all upstream regulators of the IFNy pathway, and these clusters are adjacent on the visualization). Moreover, as described above, we randomly sampled 1,000 cells expressing NT gRNA and re-labeled them as a new targeted gene class, representing a negative control (NEG CTRL). Despite receiving a different label in the LDA procedure, these cells were indistinguishable from NT controls in the resulting embedding, demonstrating that our procedure does not overfit the data (Supplementary Figure 9).

### Differential expression and gene set enrichment analyses

We used FindMarkers() in Seurat to find differentially expressed genes between non-targeting cells and cells that belonged to a targeted gene class. The top 20 genes from each class were used as input into the heatmap in Figure 4A. Finally, this top300 list of genes from each class was used as input into the EnrichR package [52,53] to run pathway analysis using the human WikiPathways database from 2019. Figure 4F shows the top five enriched pathways with a p_value < 0.001 for *CUL3* KO cells.

### *NRF2* overexpression experiments

*NRF2* over-expression plasmid was purchased from Addgene (#21549). To transfect THP-1 cells, GeneXplus reagent was used as recommended by the manufacturer. 24 hours post-transfection cells were inspected under the microscope to verify reporter eGFP and dsRed proteins were expressed in the cells. 24-48 hours post-transfection, cells were collected and washed with R10 media. Flow cytometry was used to assess changes in PD-L1 protein expression as previously described.

### JQ1 inhibitor experiments

THP-1 cells were treated with DMSO, JQ1 (1μM, 24 hours), JQ1 + IFNγ, Decitabine+TGFβ1+IFNγ or Decitabine+TGFβ1+IFNγ +JQ1. PD-L1 expression was assessed by flow cytometry as previously described.

### Validation CRISPR screen

We designed new gRNAs using the *guides* webtool to target *KEAP1, NRF2, BRD4* and *CUL3* in order to validate our ECCITE-seq findings. Plasmids containing the gRNAs were pooled at equal ng amounts and the virus was produced as previously described. THP-1 cells were transduced at low MOI and cells were selected with blasticidin for three days. After selection Cas9 expression was induced and cells were stimulated as previously described. At the end of stimulation, cells were spun down, resuspended in 100μl of MACS buffer containing 5μl of FcX blocking reagent and placed on ice for ten minutes. Next, cells were stained with a PD-L1 antibody for 30 minutes, washed with 1mL of MACS buffer and passed through a 40μM cell strainer to remove cell clumps. The Sony SH100 sorter was used to sort the top 15% of cells with the highest and lowest PD-L1 protein expression in two separate tubes containing Quick Extract buffer (Epicenter). We amplified the gRNA sequence from the isolated genomic DNA as described in [54]. Samples we sequenced with a target recovery of 1000 reads per gRNA per sample. To quantify gRNA counts in each sample, we first made a gRNA reference fasta file and used it to map and quantify our reads with Bowtie2 [55]. To analyze our data and find gRNAs enriched or depleted in our samples we used MAUDE [56].

