## Supplementary Figures for "Characterizing the molecular regulation of inhibitory immune checkpoints with multi-modal single-cell screens"

### Supplementary Figure 1

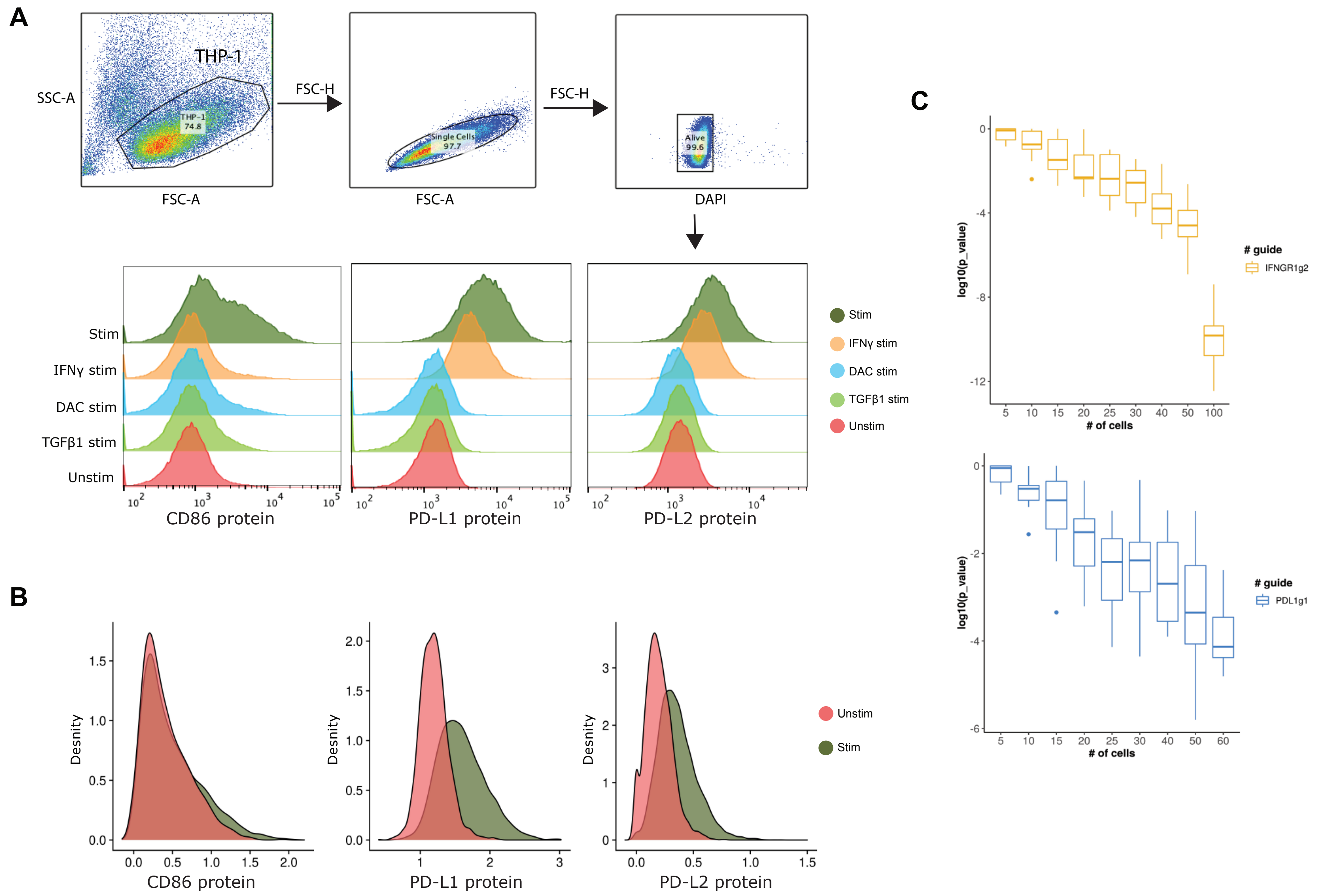


**Supplementary Figure 1. THP-1 cell stimulation induces the expression of multiple immune checkpoint proteins (related to Figure 1).**
**(A)** Top: Example gating strategy for flow cytometry to remove cell doublets and dead cells. Bottom: Single-cell expression levels for CD86, PD-L1 and PD-L2 proteins across different stimulation conditions, as measured by flow cytometry. **(B)** Single-cell expression levels for CD86, PD-L1 and PD-L2 proteins across different stimulation conditions, as measured by CITE-seq. Relative levels are concordant with (A). **(C)** Power analysis to estimate the number of cells necessary to detect statistically significant shifts in protein expression across two different gRNAs (IFNGR1g2: yellow, PDL1g1: blue). Each boxplot summarizes ten random sampling draws of the indicated number of cells and the log-transformed p-values generated through differential protein expression analysis (DE) using the Wilcoxon Rank sum test. DE was performed using the same number of sampled IFNGR1g2, PD-L1g1 and non-targeting control cells.

### Supplementary Figure 2

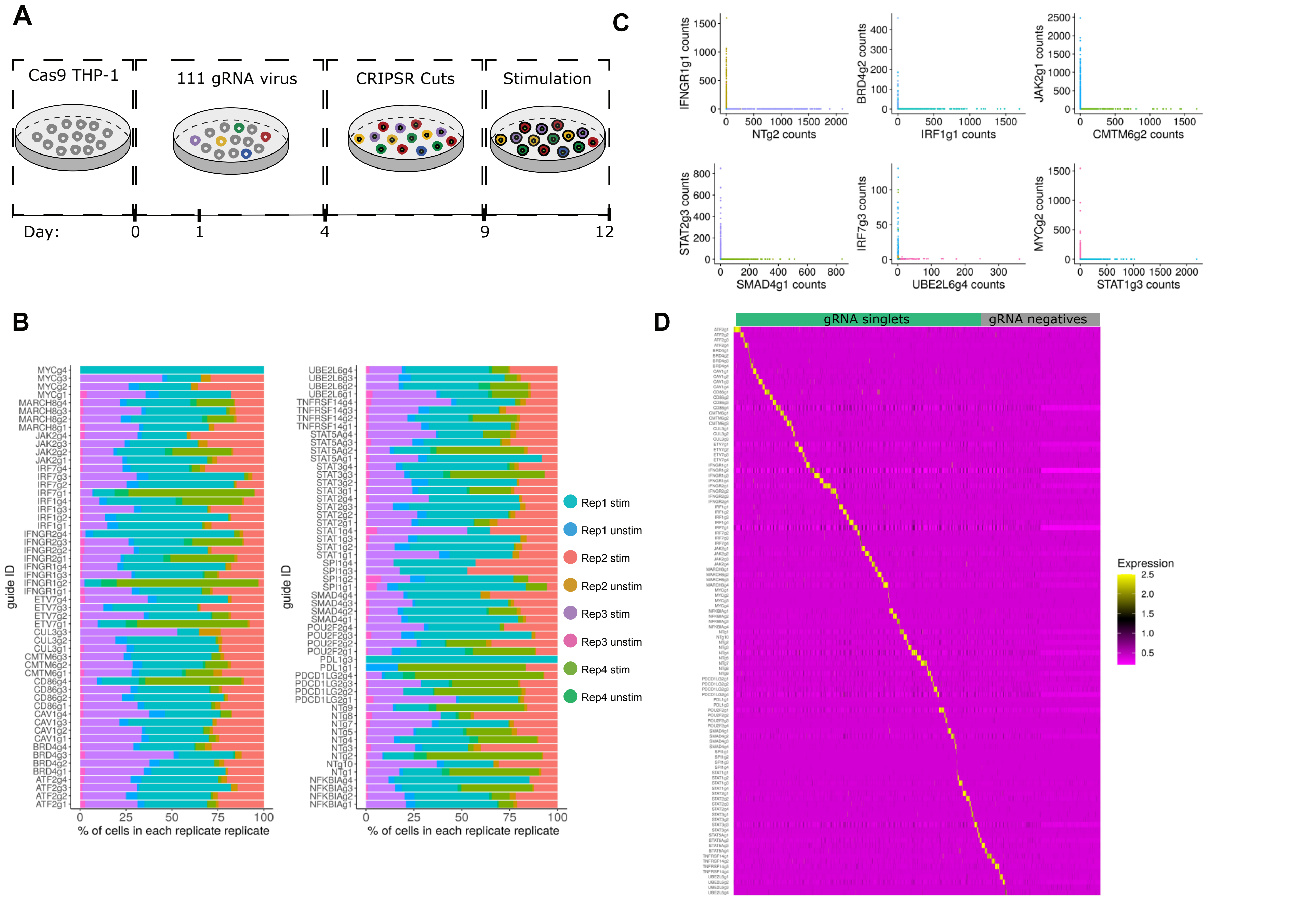


**Supplementary Figure 2. Pre-processing of the 111-gRNA ECCITE-seq experiment (related to Figure 2).**
**(A)** Schematic representation of experimental setup. A Cas9-inducible THP-1 cell line was transduced with a virus containing 111 gRNAs (low MOI). Cells were treated with Blasticidin for three days to select for successfully transduced cells. Next, cells were treated with doxycycline to induce Cas9 expression and CRISPR-mediated gene KOs for 5 days. Lastly, cells were stimulated with TGFβ1, Decitabine and IFNγ for three days as previously described. ECCITE-seq was performed 12 days post viral transduction. **(B)** Barplots showing gRNA representation for each replicate (four biological replicates, split also by stimulation condition: stim and unstim). While replicates 1-3 had even representation of gRNAs, replicate 4 (oldest transduction) had skewed gRNA representation, likely as a result of long term cell culture. Replicate 4 was excluded from downstream analyses. **(C)** Example pairwise scatter plots showing mutually exclusive gRNA expression in single cells. number of sampled IFNGR1g2, PD-L1g1 and Non-targeting control cells. **(D)** gRNA expression heatmap showing the majority of cells express only one gRNA (singlets: green, negatives: grey). We also detected 30 gRNA doublet cells that are not shown here. gRNA counts are normalized (CLR normalization) and scaled (z-score).

### Supplementary Figure 3

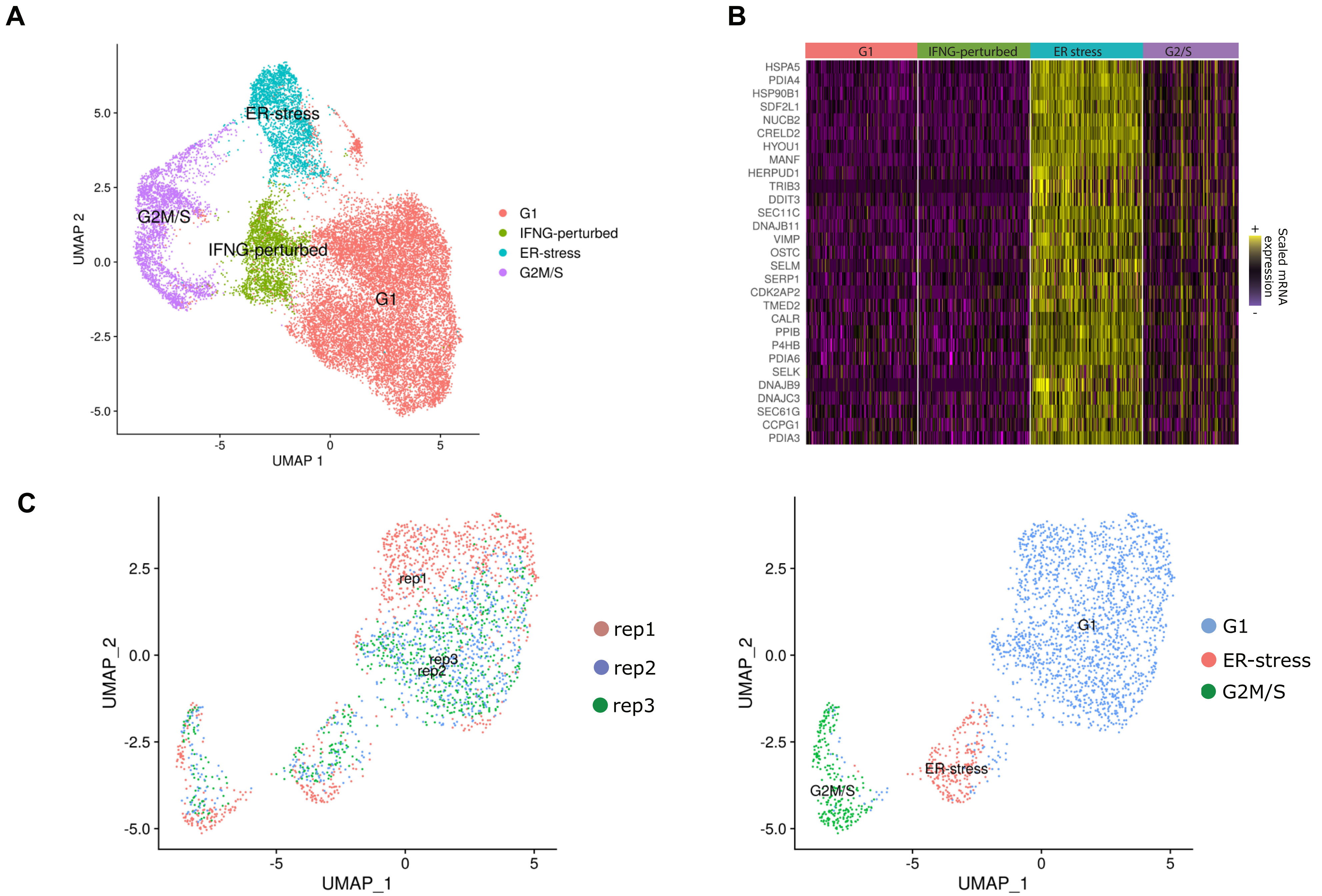


**Supplementary Figure 3. Unwanted sources of variation drive mRNA-based clustering (related to Figure 2).**
**(A)** UMAP visualization of the ECCITE-seq dataset based on cellular transcriptomes. Clusters are driven by different sources of variation shown in different colors (cell cycle state, CRISPR perturbation, stress). Figure is similar to Figure 2A, but with labels for the ER-stress cluster. **(B)** Single-cell heatmap showing the up-regulation of a specific gene module in the ER-stress cluster. EnrichR analysis demonstrates that this gene set is enriched (adjusted p-value < $5*10^{-20}$) for 'response to endoplasmic reticulum stress". **(C)** Similar to (A), but computed using only only NT cells. This demonstrates that confounding sources of heterogeneity are present even in the absence of perturbation.

### Supplementary Figure 4

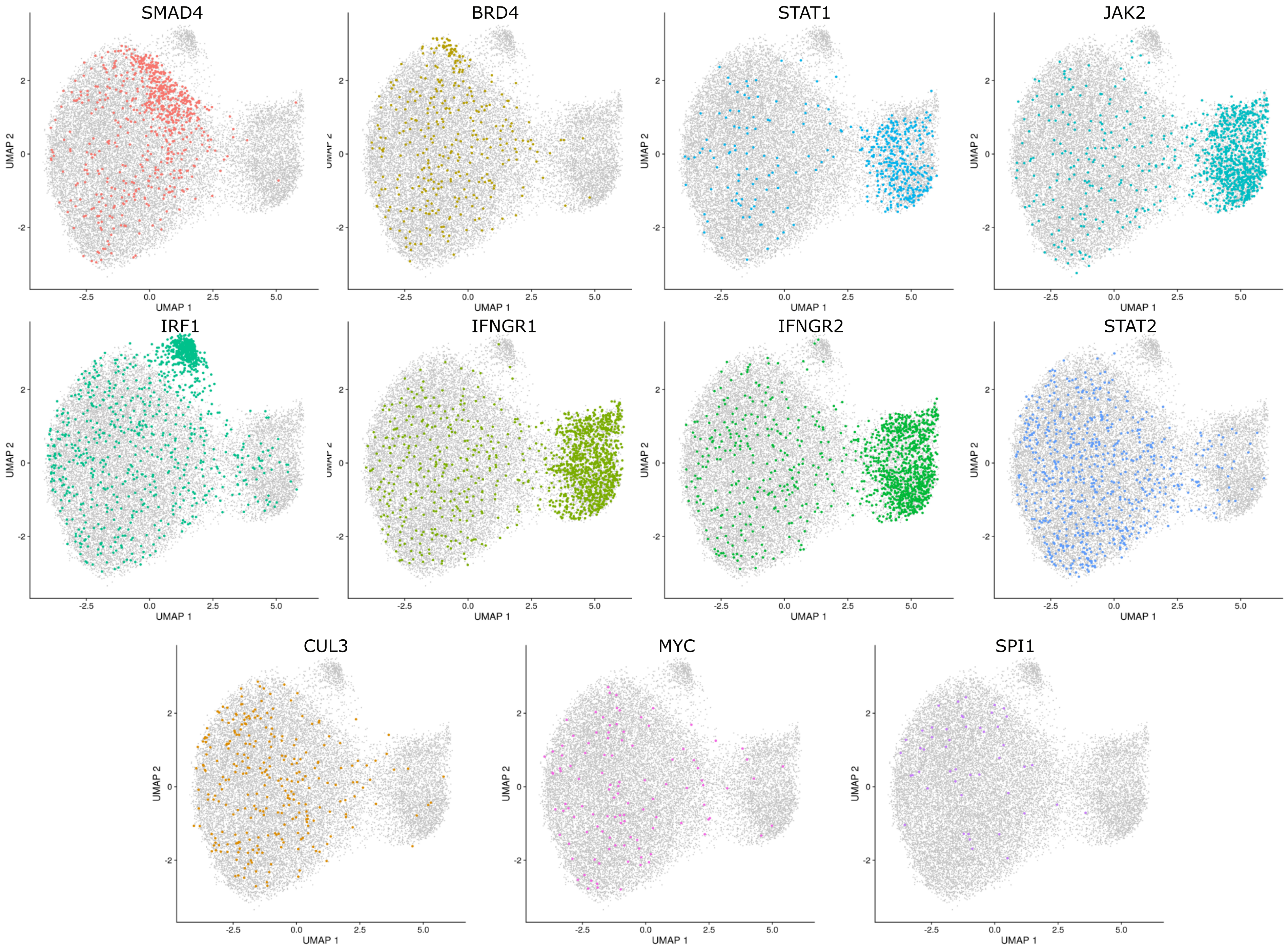


**Supplementary Figure 4. Calculating local perturbation signatures controls for unwanted sources of variation.**
Similar to Figure 2D, but the cells from each individual perturbation are specifically highlighted. In addition to some perturbations which form specific clusters (e.g. IRF1), other perturbations (e.g. BRD4 and SMAD4) exhibit weaker evidence of sub-clustering, suggesting that improved analysis strategies would help to reveal their perturbation state.

### Supplementary Figure 5

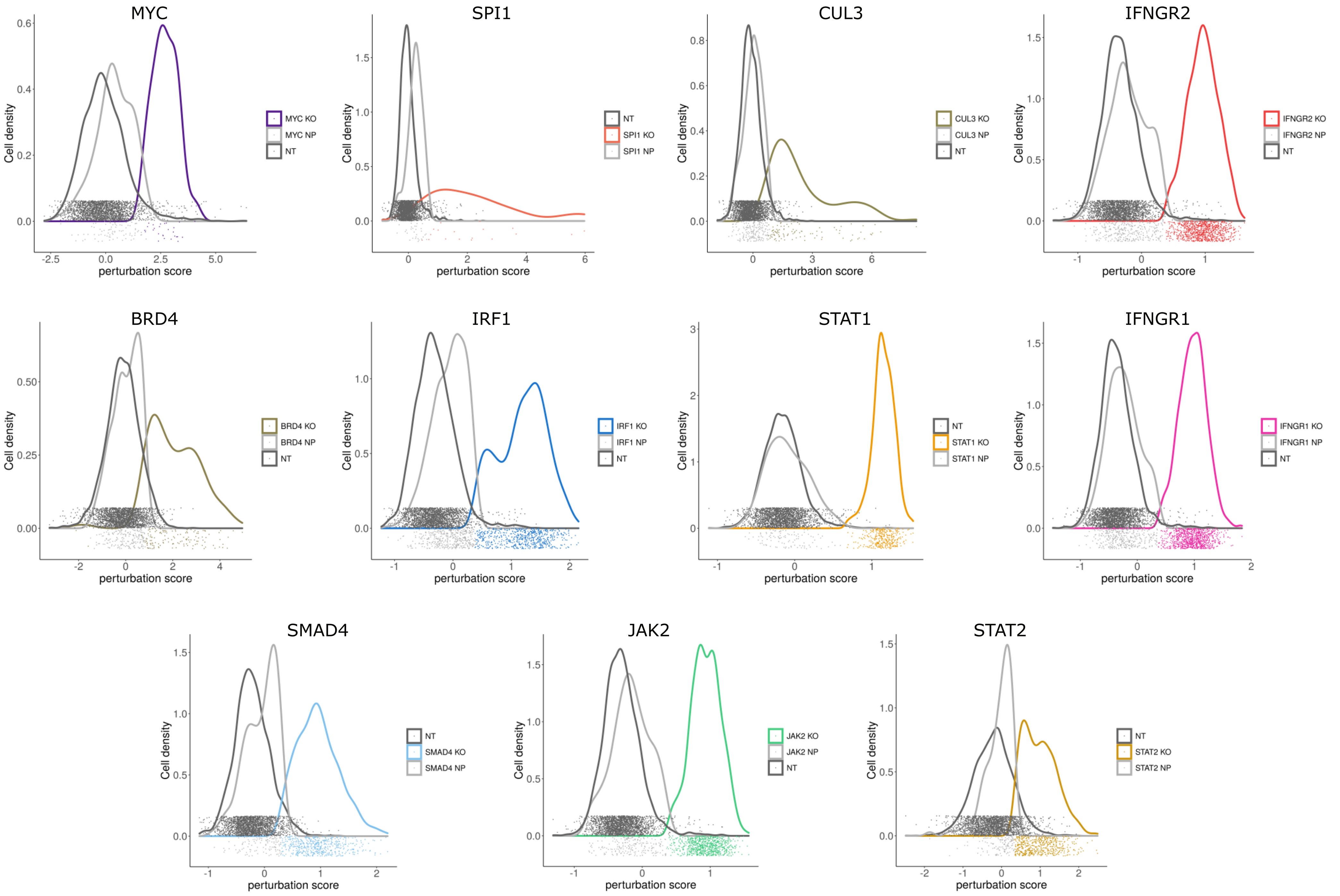


**Supplementary Figure 5. Mixscape models targeted cells as a heterogeneous mixture.**
For each cell, we calculated a perturbation score (Supplementary Methods) representing its strength of perturbation compared to the average of NT controls. We calculated this not only for targeted cells, but also for cells expressing NT gRNA in order to estimate the variance in the control population. Here, we show the distribution of perturbation scores as a function of mixscape classification (similar to Figure 3A). Dots on the x-axis represent single-cell values, and are colored to match the mixscape classifications. Non-perturbed cell densities (NP, light grey) overlap with the non-targeting control cell densities (NT, dark grey).

### Supplementary Figure 6

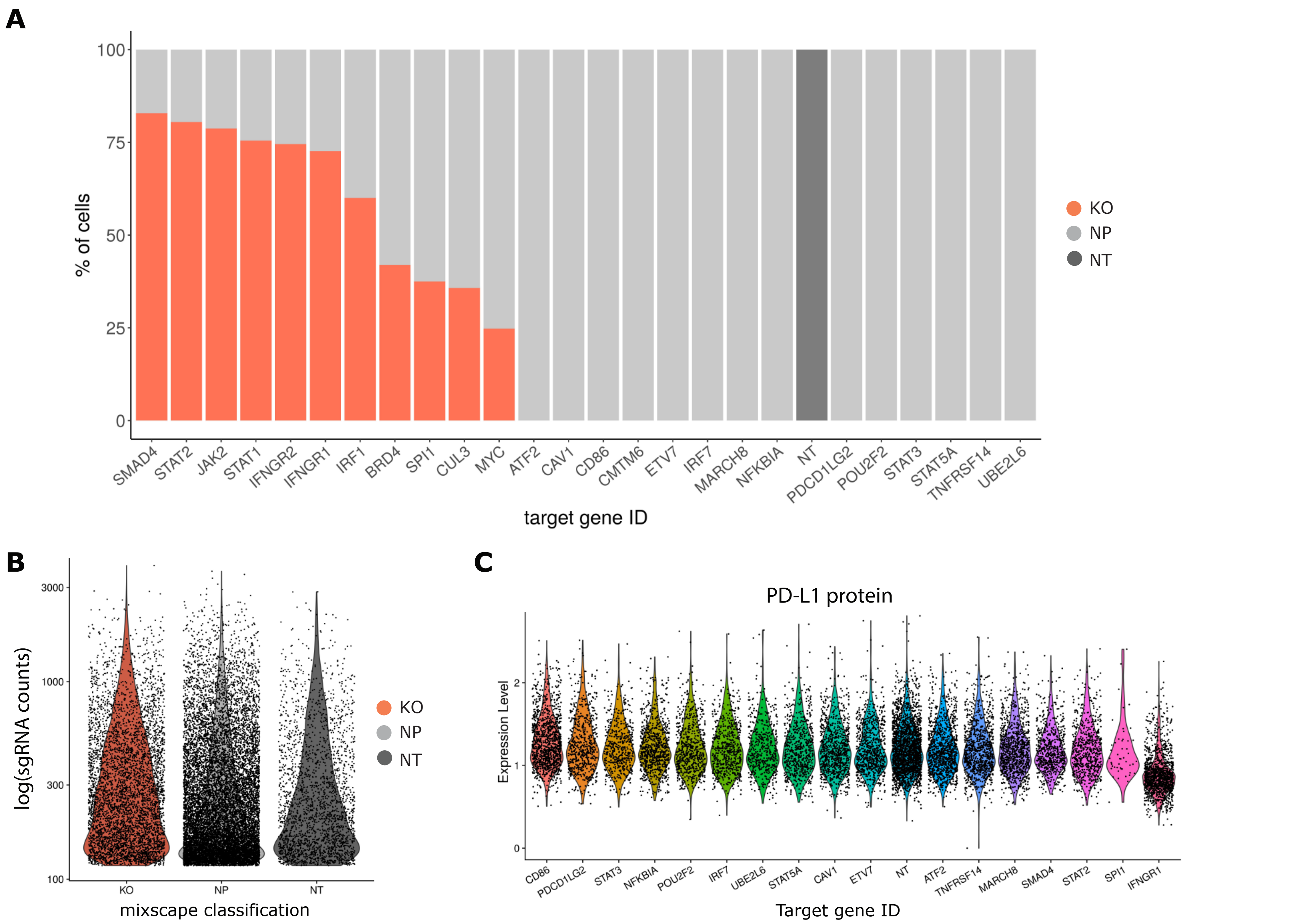


**Supplementary Figure 6. Mixscape does not identify perturbed cells for all target genes.**
**(A)** Barplots showing the % of knockout (KO, red) and non-perturbed (NP, light grey) cells as classified by mixscape within each target gene class. Some target gene classes contain only NP cells because we did not observe detectable transcriptomic shifts in any cells. **(B)** Violin plot showing the log-transformed gRNA counts in the dataset split by the global mixscape classification (NT, KO and NP). All three groups have similar count distributions suggesting that the NP cells are not a result of incorrect gRNA classification. **(C)** Violin plots showing PD-L1 protein levels in many target gene groups remain unchanged in response to their perturbation. For reference, IFNGR1 KO cells are included.

### Supplementary Figure 7

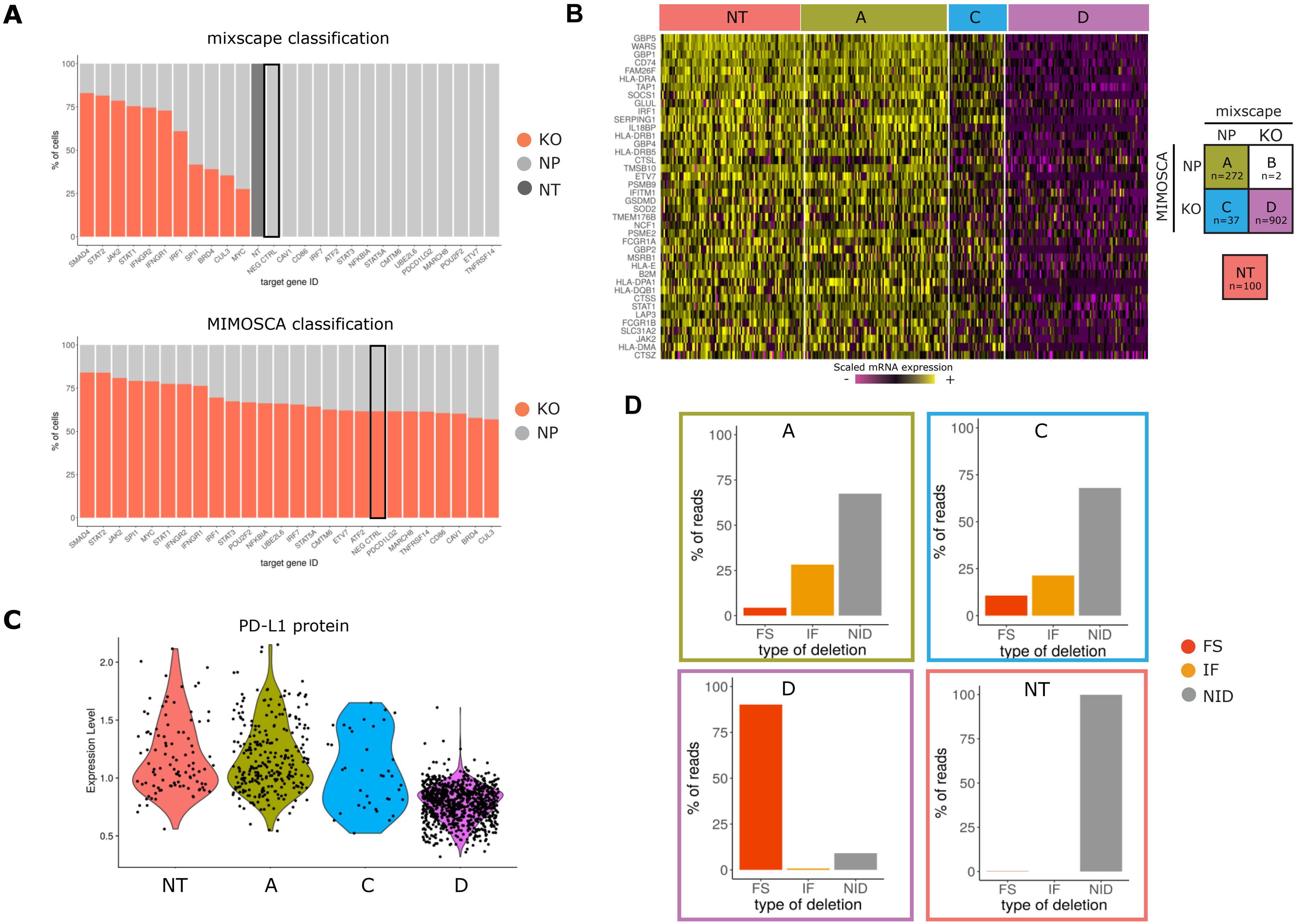


**Supplementary Figure 7. Benchmarking mixscape against MIMOSCA.**
**(A)**Top: Barplots showing the % of KO (red) and NP (light grey) cells within each gRNA class as classified by mixscape, and MIMOSCA (Bottom). To assess the potential for overfitting, prior to running the dataset, we randomly sampled 1,000 cells expressing NT gRNA and re-labeled them as a new targeted gene class, representing a negative control (Neg_CTRL, marked with a black box). Only mixscape correctly classifies all of these cells as NP. **(B)** Single-cell mRNA expression heatmap with IFNGR2g2 cells being grouped by mixscape and MIMOSCA classification. Cells classified by both methods as KO (Class 'A') exhibit downregulation of IFNγ pathway genes, while cells classified by both methods as ND (Class 'D') resemble NT controls. When mixscape classifies cells as NP and MIMOSCA classifies as KO (Class C), cells resemble NT controls, suggesting that the mixscape classification is correct. Class B (2 cells total) was removed for visualization due to low cell number. **(C)** Violin plots showing PD-L1 protein expression in IFNGR2g2 cells grouped by their MIMOSCA and mixscape classification (see legend in (B)). Class C cells resemble NT controls, suggesting that the mixscape classification is correct. **(D)** Barplot showing the % of reads with no INDELS (grey), inframe (orange) and frameshift (red) mutations across all MIMOSCA and mixscape IFNGR2g2 cell classifications. Class C cells resemble NT controls, suggesting that the mixscape classification is correct.

### Supplementary Figure 8

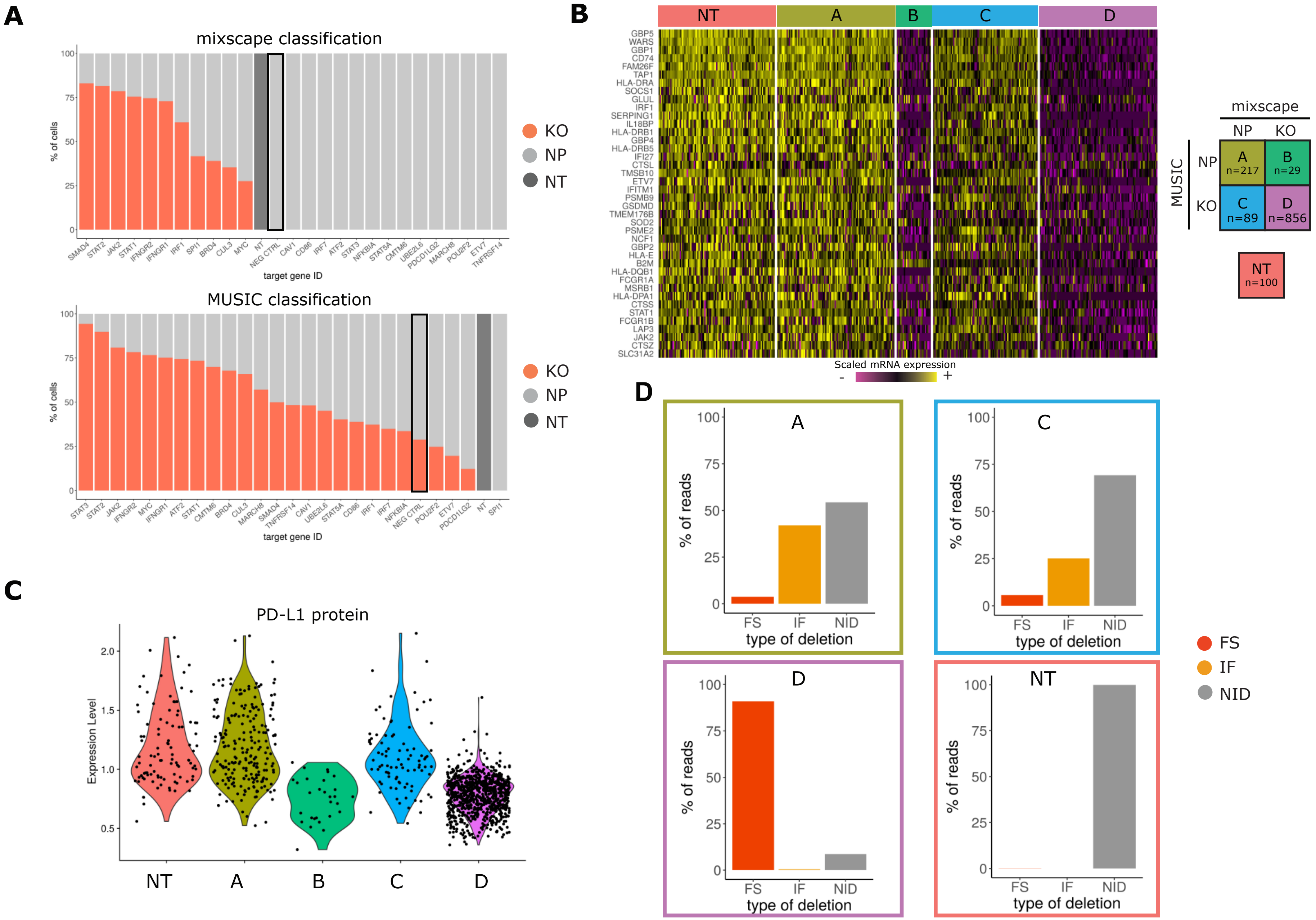


**Supplementary Figure 8. Benchmarking mixscape against MUSIC.**
**(A)** Top: Barplots showing the % of KO (red) and NP (light grey) cells within each gRNA class as classified by mixscape, and MUSIC (Bottom). To assess the potential for overfitting, prior to running the dataset, we randomly sampled 1,000 cells expressing NT gRNA and re-labeled them as a new targeted gene class, representing a negative control (Neg_CTRL, marked with a black box). Only mixscape correctly classifies all of these cells as NP. **(B)** Single-cell mRNA expression heatmap with IFNGR2g2 cells being grouped by mixscape and MUSIC classification. Cells classified by both methods as KO (Class 'A') exhibit downregulation of IFNγ pathway genes, while cells classified by both methods as ND (Class 'D') resemble NT controls. When mixscape classifies cells as NP and MUSIC classifies as KO (Class C), cells resemble NT controls. When mixscape classifies cells as KO and MUSIC classifies as NP, cells exhibit evidence of perturbation. Therefore, groups B and C suggest that when the methods disagree, the mixscape classification is correct. **(C)** Violin plots showing PD-L1 protein expression in IFNGR2g2 cells grouped by their MUSIC and mixscape classification. Groups B and C suggest that when the methods disagree, the mixscape classification is correct. **(D)** Barplot showing the % of reads with no INDELS (grey), inframe (orange) and frameshift (red) mutations across all MUSIC and mixscape IFNGR2g2 cell classifications. Groups B and C suggest that when the methods disagree, the mixscape classification is correct.

### Supplementary Figure 9

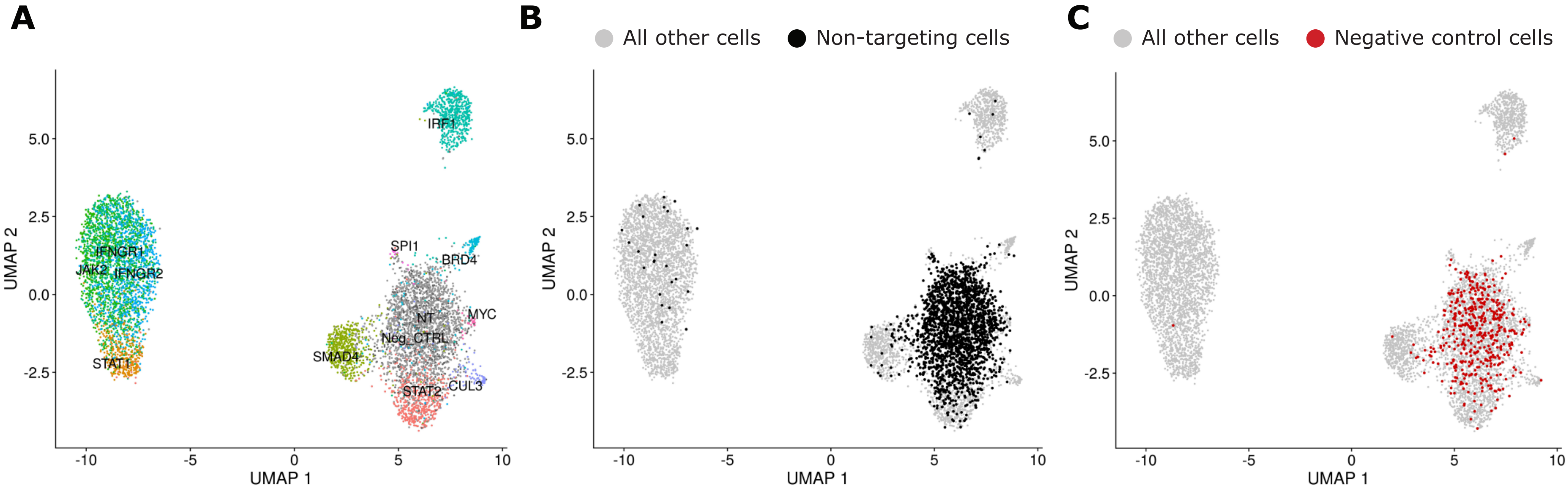


**Supplementary Figure 9. Linear Discriminant Analysis for the clustering of ECCITE-seq data (related to Figure 3).**
**(A)** UMAP visualization of all 7,421 NT, KO and negative control (Neg_CTRL) cells after running Linear Discriminant Analysis (Supplementary Methods). Cells from each targeted class separate visually on the 2D embedding. However, Neg_CTRL cells cluster with NT cells despite having a distinct label, demonstrating that our procedure does not overfit the data and induce separation when there are no transcriptomic differences. **(B)** Same as in (A), highlighting the non-targeting cells (black). **(C)** Same as in (A), highlighting the negative control cells (red).

### Supplementary Figure 10

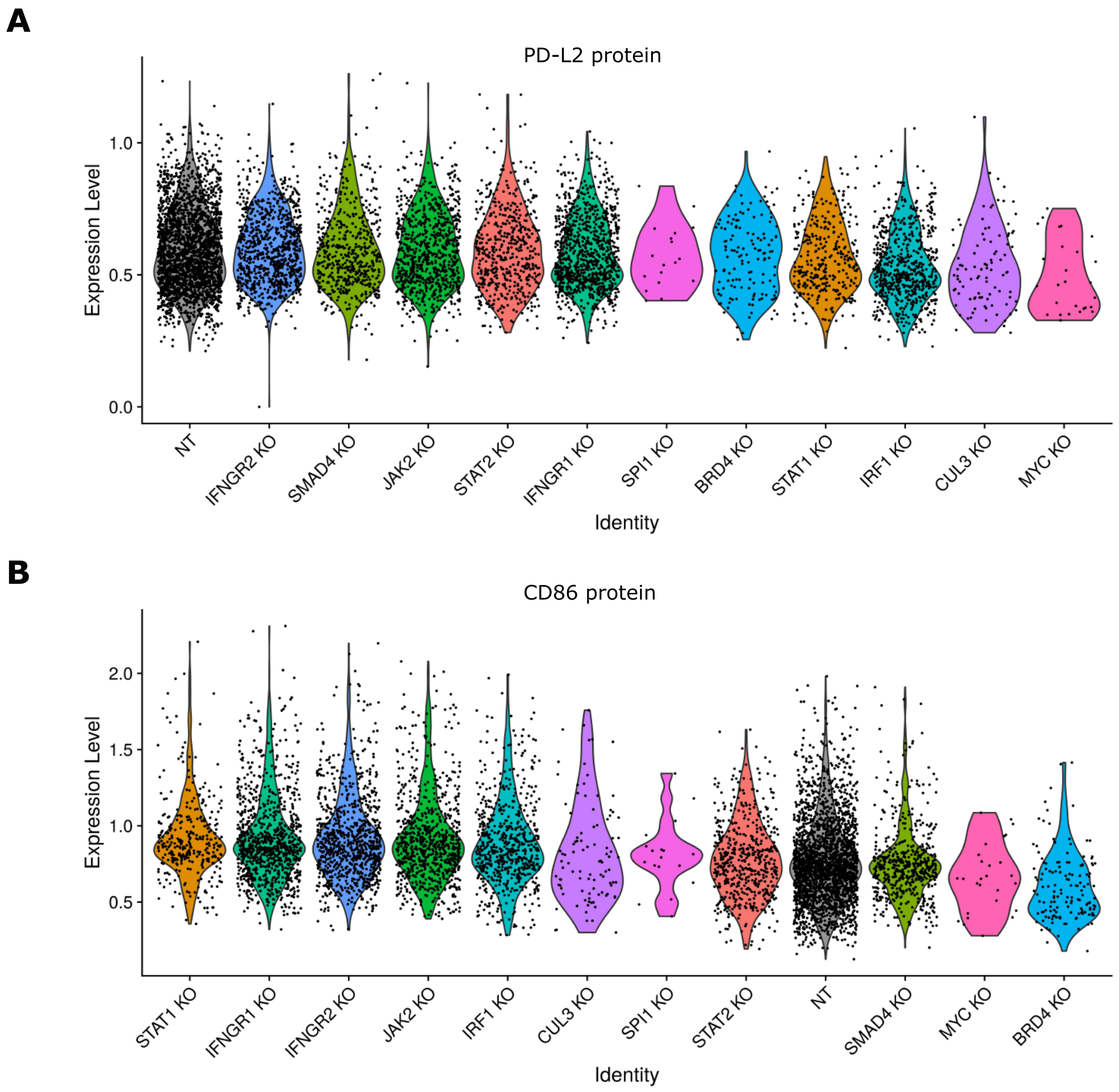


**Supplementary Figure 10. CD86 and PD-L2 expression in KO cells.**
**(A)** Violin plots showing PD-L2 protein levels across KO target gene classes. **(B)** Violin plots showing CD86 protein levels across KO target gene classes.

### Supplementary Figure 11

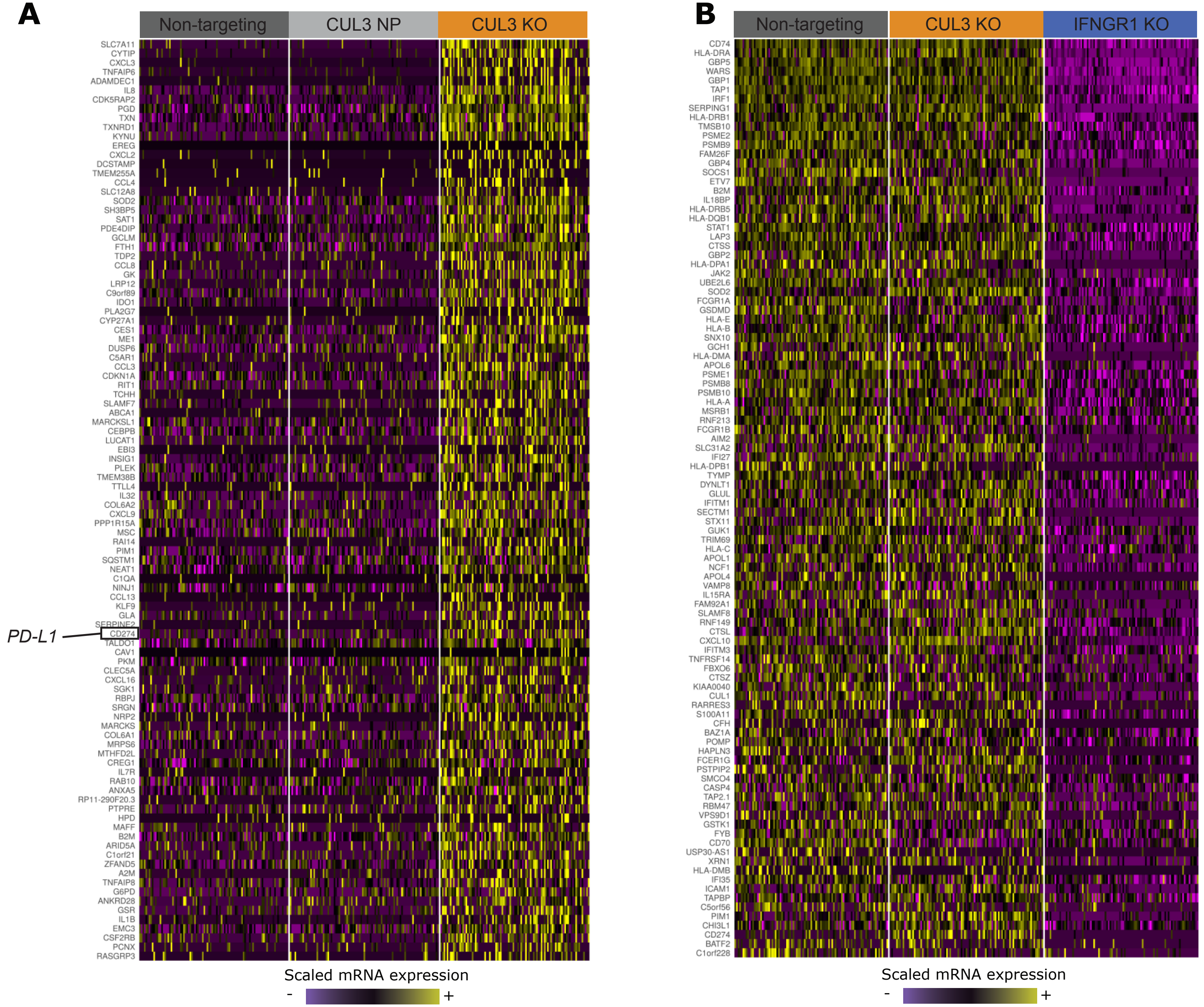


**Supplementary Figure 11. *CUL3* KO cells have a unique transcriptomic signature.**
**(A)** Single-cell mRNA expression heatmap showing that *CUL3* KO cells upregulate a module of genes in comparison to NT and *CUL3* NP cells (including the *PD-L1* transcript, *CD274*, highlighted on the heatmap). **(B)** Single-cell mRNA expression heatmap showing that the CUL3 transcriptomic signature is not IFNγ-related, suggesting *CUL3* is acting through an alternative pathway to regulate *PD-L1* at the transcriptional level. For both (B) and (C) heatmaps, lists of genes were obtained using FindMarkers() function in Seurat (Wilcoxon Rank sum test). mRNA counts are log-normalized and scaled (z-score).

### Supplementary Figure 12

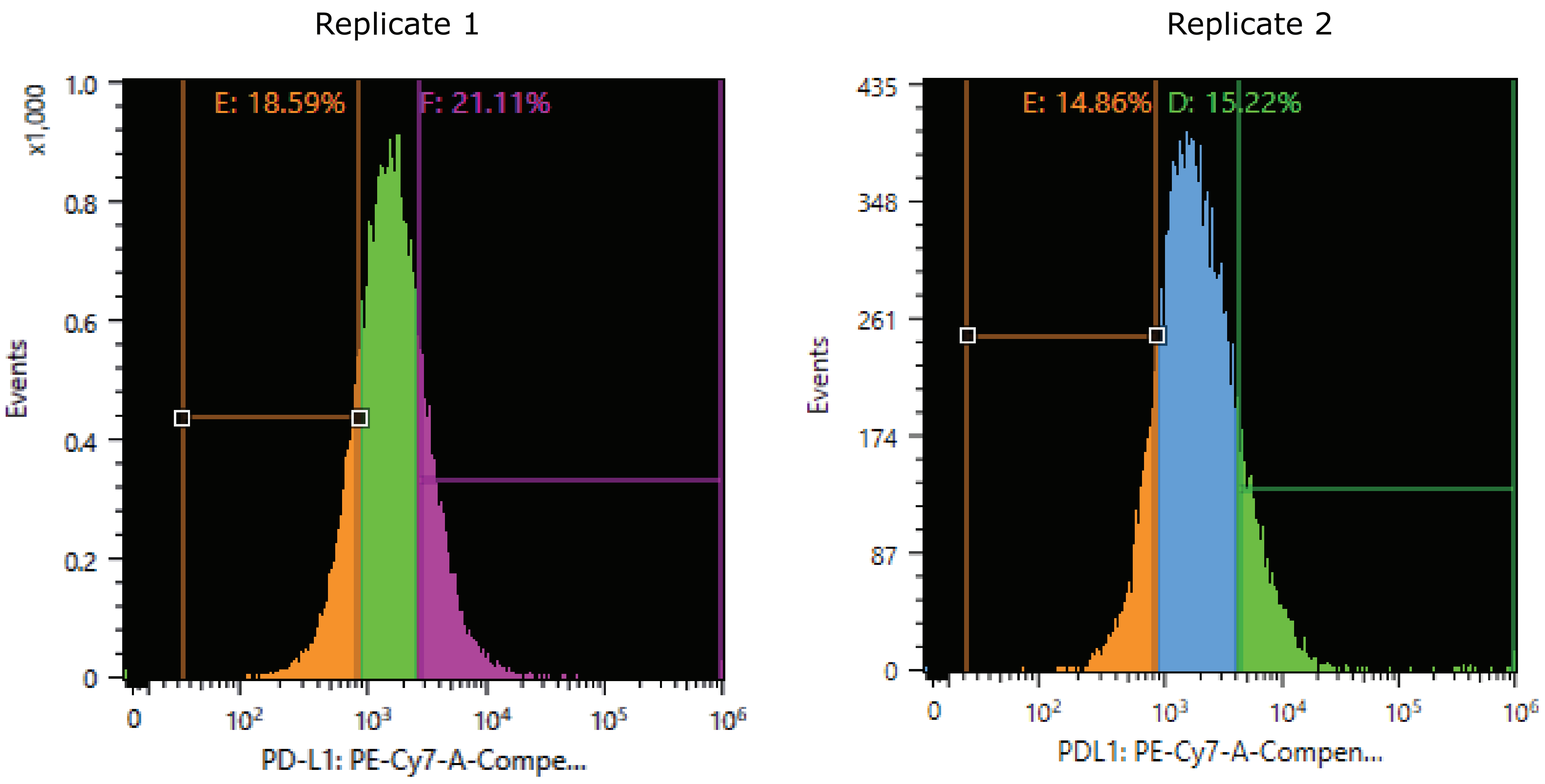


**Supplementary Figure 12. FACS gating strategy for validation pooled CRISPR screen (related to Figure 5).**
Flow cytometry plots showing gating for PD-L1 high and PD-L1 low expressing cells. Each gate represents 15-20% of the total number of cells in the data. Gating strategy is shown for both replicates.
